# Connectivity Regression

**DOI:** 10.1101/2023.11.14.567081

**Authors:** Neel Desai, Veera Baladandayuthapani, Russell T. Shinohara, Jeffrey S. Morris

## Abstract

Assessing how brain functional connectivity networks vary across individuals promises to uncover important scientific questions such as patterns of healthy brain aging through the lifespan or dysconnectivity associated with disease. In this article we introduce a general regression framework, Connectivity Regression (*ConnReg*), for regressing subject-specific functional connectivity networks on covariates while accounting for within-network inter-edge dependence. ConnReg utilizes a multivariate generalization of Fisher’s transformation to project network objects into an alternative space where Gaussian assumptions are justified and positive semidefinite constraints are automatically satisfied. Penalized multivariate regression is fit in the transformed space to simultaneously induce sparsity in regression coefficients and in covariance elements, which capture within network inter-edge dependence. We use permutation tests to perform multiplicity-adjusted inference to identify covariates associated with connectivity, and stability selection scores to identify network edges that vary with selected covariates. Simulation studies validate the inferential properties of our proposed method and demonstrate how estimating and accounting for within-network inter-edge dependence leads to more efficient estimation, more powerful inference, and more accurate selection of covariate-dependent network edges. We apply ConnReg to the Human Connectome Project Young Adult study, revealing insights into how connectivity varies with language processing covariates and structural brain features.

## 1. Introduction

Functional connectivity, the study of temporally coincident brain activity, seeks to link geographically distinct brain regions via measures of statistical dependence (Toga, 2015). Since the brain is known to sub-specialize regions for distinct responsibilities, functional connectivity sheds insight into which brain regions coordinate together by assessing mutual brain activation measured by blood oxygenation scores (BOLD) from functional MRI (fMRI) images across timepoints (Eickhoff et al., 2015). fMRI is typically classified as task-based or resting state; while task-based fMRI seeks to identify brain-regions involved in a particular activity, resting state fMRI explores the intrinsic sub-specialization of brain regions at rest and how they organize together for core cognitive functions (Zhang et al., 2016; Lv et al., 2018). In practice, functional connectivity is often represented as a network/graph based on a correlation, covariance, or precision matrix, with an entry denoting dependencies between brain regions assessed from BOLD scores (Stulz et al., 2018; Mohanty et al., 2020). For functional connectivity derived from resting state fMRI, active research areas involve either the accurate estimation of these networks or an assessment of how functional connectivity varies with covariates in order to determine the biological link between phenotype and the brain’s natural functional organization (Zhu et al., 2019). The core scientific aim of this paper is the latter; we seek to develop a rigorous statistical framework to link phenotype to functional connectivity networks derived from resting state fMRI.

The Human Connectome Project (HCP) aimed to construct a comprehensive network mapping the human brain to provide insight into anatomic and functional connectivity and aid research into disorders such as autism, Alzheimer’s, and schizophrenia. Using a variety of non-invasive imaging methods, the project collects extensive connectome information for healthy, diseased, and longitudinally sampled patients. Among its large scale goals to comprehensively map brain circuits, a major application of the HCP is to assess the link between patient connectivity structure and phenotypic traits (Van Essen et al., 2013). The link between brain connectivity and phenotype has the potential to provide deeper nuanced insight into brain function. For example, contrasting overall connectivity of diseased and healthy populations highlights circuitry changes potentially driving diseases (Zhao et al., 2020). Analogously, a comparison of connectivity structure across ages helps scientists understand the biological underpinnings of the maturation process (Lawrence et al., 2019). While these examples compare global connectivity differences, a more ambitious goal is to link phenotypic changes explicitly with circuits between specific brain regions. Given rapid advances in neuroscientific knowledge, finding precise region-specific links with phenotype can better integrate existing biological knowledge and determine the relationship between structure and function. To this end, in this paper we develop a general regression approach, Connectivity Regression (*ConnReg*), that relates subject-specific network connectivity objects to a set of subject-specific covariates and then characterizes which network circuits are driving these differences. We begin with a simple example in Figure 1 to illustrate this goal and to describe one of the key conceptual underpinnings of our approach: inter-edge association that engenders a type of second order dependence.

**Figure 1.**
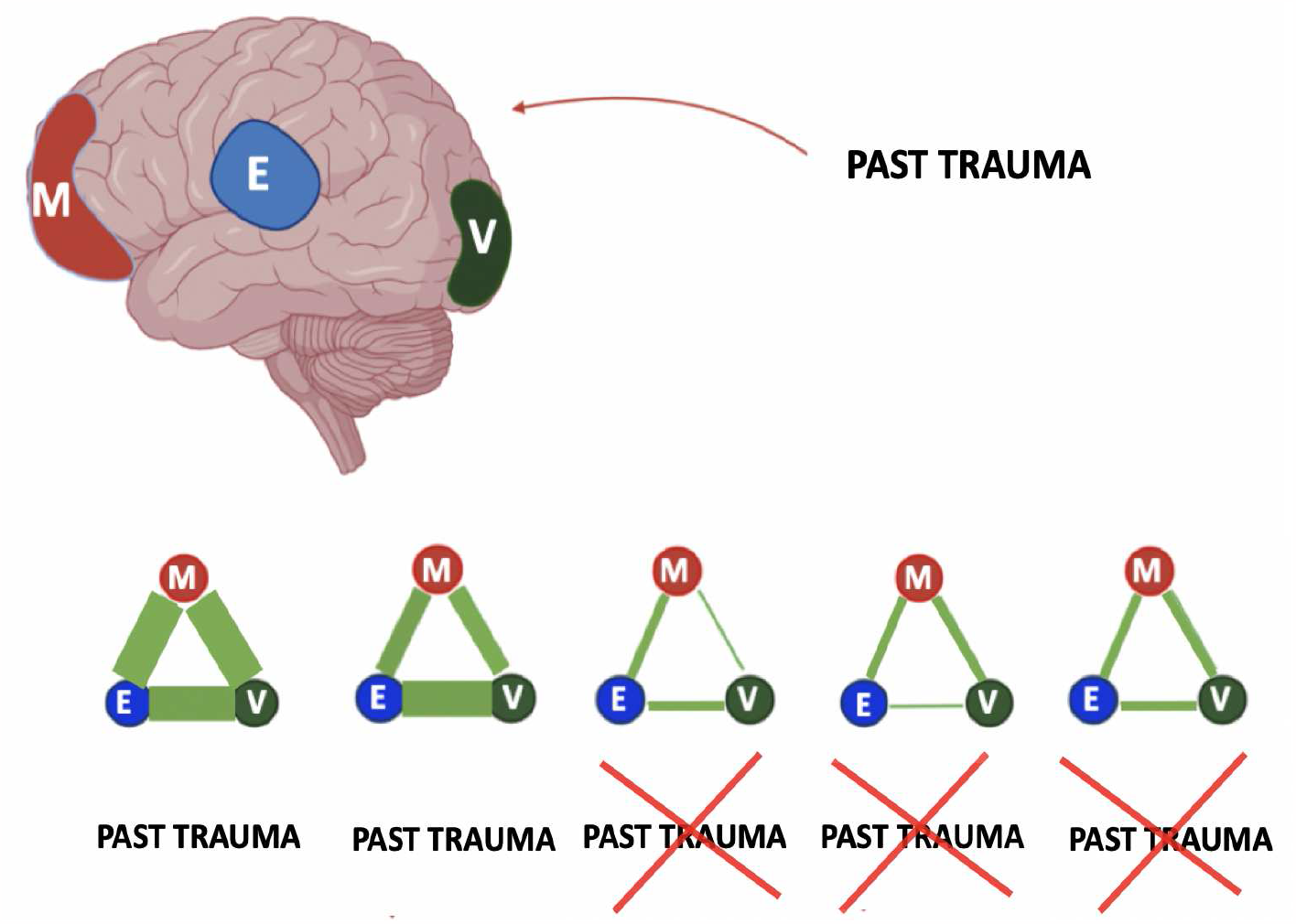
Conceptual overview: We divide the brain into 3 regions: vision (V), memory (M), and emotion (E) centers. The middle panel represents subject-specific connectivity networks for four individuals, with line thickness indicating the network circuit’s strength of association. The covariate considered is a dichotomous indicator for whether an individual suffered a past trauma. The core goal of connectivity regression is to assess if covariates affect connectivity objects. In this case “past trauma” is associated with connectivity, affecting all three circuits. Independent of “past trauma”, the magnitude of the vision and memory circuit, the memory and emotion circuit, and the vision and emotion circuit move up and down in concert with each other. This demonstrates the concept of inter-edge dependence, a type of second order association that we account for in our modeling.

Motivated by functional data analysis (FDA) approaches, we undertake a joint modeling approach that accounts for the internal structure of the network object. In the setting of FDA, the internal structure is the within-function correlation structure of the functional data object *Y* (*t*) in that observations at nearby domain points *t* are similar to one another; models incorporating smoothness in *t* and accounting for correlation in the between-function residuals lead to more efficient estimation and more powerful joint inference (Morris, 2015). In a similar spirit, for these connectivity network objects the internal structure involves inter-edge dependence structures, as illustrated in Figure 1. In the toy example, note that individuals with strong connectivity in their vision-memory circuits also have strong connectivity in their memory-emotion and vision-emotion circuits (and vice-versa). This suggests that these circuits operate in concert with each other, inducing an association among the corresponding edges, a type of “correlation of correlations”. We wish to estimate and account for this type of second order association in our modeling that, as we will demonstrate, tends to produce more efficient estimation and inference and also reveals additional insights into the connectivity dynamics.

### Network-Object Regression: Regression approaches treating entire subject-specific networks as units of observation

In contrast to our approach, most preceding network regression approaches assume subject-specific vectors as their units of observation. Many of these methods focus on differences in network structure between groups, or other discrete covariates (Xia et al., 2015; Liu et al., 2017; Peterson et al., 2015; Durante et al., 2018). Ni et al. (2021) has published a review paper on Bayesian graphical models for biomedical applications which covers some methods on group graphs and their applications. The graphical model literature has also produced methods for the general regression of network edges for observed binary data (Cheng et al., 2014), directed acyclic graphs (Ni et al., 2019), and undirected graphs (Wang et al., 2021). For regression of covariance matrices on covariates, much work assumes a natural ordering of the multivariate dimensions, including autoregressive conditionally heteroscedastic models that take sequential fits of covariance across time series (Engle and Kroner, 1995) and Cholesky-decomposition based models (Pourahmadi et al., 2007). Among methods for unordered variates, Chiu et al. (1996) performed joint regression of the mean and covariance on a multivariate vector, utilizing the matrix log-transformation to model covariance elements. Hoff and Niu (2012) took a random effects approach and parameterized heteroscedastic covariance by the inner product of random effect regression coefficients.

In our setting, we have sufficient data from each subject such that we are able to obtain an entire *K* ×*K* subject-specific connectivity network across *K* brain regions as opposed to a *K*-dimensional vector. There are a handful of previous papers in the literature modeling subject-specific networks as units of observation, making various tradeoffs among interpretability, scalability, and modeling higher order structure like inter-edge dependence that can be leveraged in this setting. Seiler and Holmes (2017) focused on covariance matrices and introduced two models. Both models implicitly assumed a Gaussian distribution on observed vectors and restricted their attention to discrete covariates representing group identities; these models also do not seek to directly model replicated network edges. Zhao et al. (2020) proposed a two-step procedure where group-ICA was applied to entire functional connectivity networks followed by a covariate-assisted principal component procedure. The authors’ model identifies global changes in functional connectivity based on covariates and mitigates multiplicity issues via the use of common linear projections; their approach does not focus on producing interpretable results for assessing network edge differences in the original space of networks. Xia et al. (2020) describe a modeling framework that takes community structure amongst edges into account to potentially improve inference, produces an interpretable result, and is solved efficiently via a convex objective function. However, the method does not seek to directly examine the effects of covariates on edges and focuses on the specific case where sub-networks are known *a priori* ; inter-edge dependence is treated as fixed and not estimated as an unknown parameter. Finally, Tomlinson et al. (2022) proposed a regression framework that regresses dissimilarity summary statistics between subject-specific networks on continuous and discrete traits and develop a variety of statistical hypothesis tests to accompany their regression approach. While the method provides a framework to determine a covariate’s impact on differences between subject specific networks, the authors do not focus on assessing inter-edge dependence or characterizing individual edge effects.

In this article we propose a new framework for regressing functional connectivity networks, represented by correlation-matrix objects, on covariates. Our approach first projects subject-specific correlation matrices to Fisher correlations using a matrix logarithm transformation which theoretically justifies Gaussianity while ensuring positive semi-definiteness (Archakov and Hansen, 2021). Using a multivariate regression approach with specification of sparsity-inducing penalties on both regression coefficients and off-diagonal precision matrix elements, we then regress connectivity on predictors while estimating and accounting for inter-edge dependence across network edges. We show this strategy leads to greater efficiency and statistical power based on the principles of seemingly unrelated regression. Furthermore, we determine which covariates affect connectivity using multiplicity-adjusted permutation tests that control experiment-wise error rates and then for selected covariates rank the edges in importance of driving these differences using a false discovery rate guided stability selection. Through extensive simulation studies we validate the performance of our permutation test and provide empirical evidence that desired edge-selection operating characteristics can be controlled via appropriately calibrated stability-selection thresholds. We apply our framework to functional connectivity network data derived from HCP (Van Essen et al., 2013). Our results demonstrate that the key findings in the transformed Fisher space comprise the dominant results in the original correlation space, and thus are interpretable in practice. We find, along with the substantial presence of second order dependence, language processing ability and two key anatomical areas to be significantly associated with changes in functional connectivity. The rest of the article is organized as follows. Section 2 describes ConnReg statistical framework and inferential procedures. Section 3 summarizes the simulation results aimed to assess the performance of ConnReg against competing methods. Section 4 shows the results of our framework applied to data from HCP and Section 5 concludes the article with a discussion of the properties and scope of our proposed framework.

## 2. Connectivity Regression

### Overview and notations

For each subject *i* = 1, …, *n*, we have a set of *p* covariates ***X***_i_ = {*X*_*ij*_, *j* = 1, …, *p*} and *K* × *K* network matrix ***R***_i_ with off-diagonal elements *R*_*ikk*_′ providing some measure summarizing functional connectivity between brain region *k* and *k*^′^ for a prespecified set of *K* brain regions. Our goal is to build a general regression framework for relating the network object ***R***_i_ to covariates ***X***_i_, identifying which covariates are associated with the connectivity network, and then characterizing which network circuits vary by each covariate. Although there are various options for choice of connectivity measure, here we focus on correlation, so that ***R***_i_ is a *K* × *K* subject-specific correlation matrix. A common approach is to perform separate regressions for each pairwise association *R*_*ikk*_′ (Scheinost et al., 2015; Zhang et al., 2016). To effectively achieve this goal, Fisher’s transformation 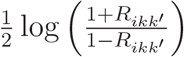 is often used to transform the pairwise correlations to the real line and engender Gaussian assumptions in the regression analyses (Fisher, 1915). This approach is insufficient for our setting since we are interested in performing a joint regression on the entire network object that accounts for its internal structure and constraints. For correlation matrix network objects, the constraints require ***R***_i_ to be a symmetric positive semi-definite matrix with 1’s down the diagonal and pairwise correlations *R*_*ikk*_′ ∈ [−1, 1] in the off-diagonals.

The internal structure includes *inter-edge dependencies*, higher order associations among the off-diagonal elements of ***R***_i_ indicating which network circuits tend to be associated with each other. Subject-specific network objects enable this type of dependence to be estimated and taken into account.

For our setting, we propose utilizing a multivariate generalization of Fisher’s transformation, the matrix logarithm transformation applied to correlation matrices. We transform the <u>entire</u> *M* = *K* × (*K* − 1)*/*2 dimensional vector of correlations ***ρ***_i_ = vecl (*R*_*ikk*_′), with vecl(·) the lower diagonal region of a symmetric matrix, into an M-dimensional vector of *Fisher correlations* ***z***_i_ = [*z*_*i*1_, …, *z*_*iM*_]^′^ ∈ ℛ ^*M*^, with each real vector mapping back to a legitimate correlation matrix. We will specify our regression model *E*(***z***|***X***) in the *Fisher space*, which we detail next, with a fast inverse mapping producing a unique and valid predicted correlation network object ***R*** for any given ***X***.

#### Fisher space transformation

The *matrix exponential* of a matrix ***R***_i_ is given by 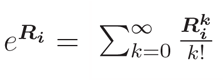(Higham, 2008). When ***R***_i_ is a *K* × *K* positive semi-definite matrix, this can be written in terms of spectral decomposition 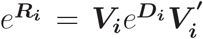, with ***D***_***i***_ a diagonal matrix of eigenvalues *λ*_*i*1_, …, *λ*_*iK*_ for ***R***_i_ corresponding to set of eigenvectors {***V***_ik_, *k* = 1, …, *K*}. In the positive semi-definite case, for a given matrix ***R***_i_, there is corresponding matrix logarithm function *logm* given by:

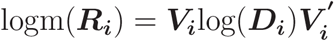

with log(***D***_i_) = {log(*λ*_1_), …, log(*λ*_*K*_)}. Archakov and Hansen (2021) introduced an algorithm for computing a matrix logarithm transformation for correlation matrices, with ***Z***_*i*_ = logm(***R***_i_) being a *K* × *K* matrix with lower triangular elements ***z***_*i*_ = vecl(***Z***_*i*_) ∈ ℛ ^*M*^, *M* = *K*(*K* − 1)*/*2. When *K* = 2, ***z***_*i*_ is a scalar and identical to Fisher’s transform. As a result, *the matrix logarithm transform of a correlation matrix can be viewed as a multivariate generalization of Fisher’s transformation*, and so we refer to the transformed space as the *Fisher space* and the elements of ***z***_*i*_ as *Fisher correlations* or *Fisher edges*.

Fisher correlations have several key beneficial modeling properties. First, there is a 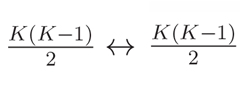 mapping between ***R*** and ***z***, with every correlation matrix ***R*** generating a unique vector ***z*** of Fisher correlations, and every real vector ***z*** ∈ ℛ ^*M*^ inducing a unique symmetric positive semidefinite *K* × *K* correlation matrix ***R*** via the efficient exponentially fast inverse mapping algorithm ***R*** = *g*^−1^(***z***) (Archakov and Hansen, 2021). This allows us to fit unconstrained regression models in the Fisher space that induce valid predicted correlation matrices. Second, unlike the Cholesky transformation, this transform is invariant to the ordering of the elements, so can be used with unordered categories. Third, the Fisher correlations are asymptotically Gaussian, so the use of normal-likelihood based models is justified in this space; justifiable Gaussian assumptions enable within network inter-edge associations to be fully captured parameterically by a covariance or precision matrix. Based on these properties, our approach is to specify a multivariate Gaussian regression model *E*(***z***|***X***) in the Fisher space that induces for each ***X*** a predicted positive semidefinite correlation matrix ***R*** = *g*^−1^(***z***).

Sparse Fisher space multivariate regression model: Given an *M* -dimensional vector of Fisher correlations ***z***_*i*_ for each subject *i* = 1, …, *n*, we assume the following multivariate regression model:

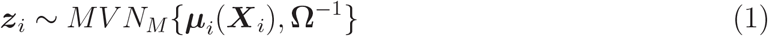

Here we assume linear regression and define the regression as ***µ***_*i*_(***X***_*i*_) = E(***z***_*i*_|***X***_*i*_) = ***β***_0_ + ***βX***_*i*_, with ***β***_0_ an *M* -vector of intercepts, ***β*** = {***β***_1_, …, ***β***_*p*_} an *M* × *p* matrix of regression coefficients corresponding to *p*-dimensional covariate vector ***X***_*i*_. The regression coefficient *β*_*mj*_ indicates the partial linear effect of covariate *X*_*j*_ on Fisher edge *m*. Predicted connectivity networks (correlation matrices) for any given ***X*** = ***x*** can be obtained by ***R*** = *g*^−1^{***µ***(***X***_*i*_ = ***x***)}. While it is valid to constrain Ω diagonal and fit *M* parallel single response models, we are primarily interested in a model allowing general Ω that can estimate and account for inter-edge dependence within connectivity networks, which as we will show, can lead to substantially more efficient estimation and inference as well as provide additional biological insights into the dynamics of brain connectivity networks. The off-diagonal elements of Ω contain partial correlations among the transformed Fisher correlation edges, in some sense a *correlation of correlations*. This allows our model to capture overdispersion in the subject-specific connectivity network circuits, introducing inter-edge dependence to account for network circuits that move in concert with one another.

We refer to this concept as *second order dependence*, since it involves the dependence among connectivity measures that are themselves measures of dependencies. Thus, multivariate regression with general Ω enables us to estimate and account for this second-order dependence in the networks.

### Motivating sparsity - connection to seemingly unrelated regressions

Zellner (1963) introduced the concept of Seemingly Unrelated Regressions (SUR), where *M* separate regression equations 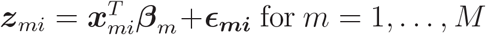 for *m* = 1, …, *M* and observations *i* = 1, …, *n* have correlated errors ***ǫ***_mi_. While each regression could validly be fit separately, accounting for residual dependence in estimation led to increased efficiency in coefficient estimation in the case where each regression had a non-identical set of predictors. In our setting, we start with the same set of predictors ***x*** across *M* and utilize a sparsity prior on ***β*** to induce different predictor sets across *M* to satisfy the conditions necessary for potential gains in estimation efficiency. For Ω, we similarly utilize a sparsity prior to both enable meaningful potential biological interpretations and to ease computational burdens associated with estimating a high dimensional object {dim(Ω) = *M* (*M* − 1)*/*2 = *O*(*K*^4^))}.

Recent literature has focused on the joint estimation of sparse coefficient matrix ***β*** and precision matrix Ω in the context of multivariate regression. These methods have been in both the Bayesian paradigm via hierachical models with a hyper-inverse Wishart prior on the covariance matrix (Bhadra and Mallick, 2013; Consonni et al., 2017) and in a penalized optimization framework (Rothman et al., 2010; Lee and Liu, 2012; Yin and Li, 2013; Cai et al., 2013). In this article we utilize the multivariate spike-slab lasso (mSSL), a penalized optimization approach that reduces to solving a series of penalized likelihood problems with a penalty that adapts until convergence (Deshpande et al., 2019). Analogous to the well known Lasso’s connection to a posteriori optimization with a Laplace prior, the mSSL has a connection to a posteriori optimization with separate priors that each consist of a mixture of Laplace distributions on coefficients and precision elements. The method is fit with an Expectation Conditional Maximization (ECM) algorithm, producing sparse point estimates of both ***β*** and Ω that have empirically been shown to produce results with excellent estimation accuracy. We next outline ConnReg_*mSSL*_, an implementation of our ConnReg framework in which we use the mSSL to induce sparsity in both ***β*** and Ω while accounting for second order dependence. Given that we use an optimization-based approach that produces sparse point estimates, we describe how we build an inferential procedure that obtains multiplicity adjusted p-values for which covariates affect functional connectivity networks and produces bootstrap-based stability selection scores to characterize which network edges are driving those effects.

### Inducing sparsity via the multivariate spike-slab lasso

Conceptually, the mSSL’s penalized multivariate optimization framework equates to placing separate spike-slab priors on the elements of ***β*** and Ω (Deshpande et al., 2019). The spike-slab lasso prior is a mixture of two Laplace distributions with a large and small scale parameter:

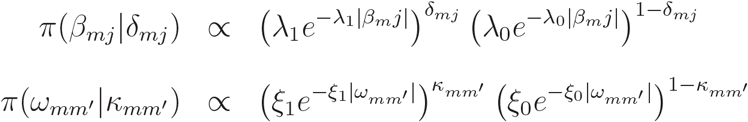

For ***β***_*mj*_, large scale parameter 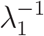 is designed to capture substantially large coefficients and small scale parameter 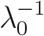 should capture negligible ones, with *δ*_*mj*_ a 0-1 indicator of whether the regression coefficient for covariate *j* and Fisher edge *m* is substantially large to be included in the model. For precision elements *ω*_*mm*_′ of Ω, the prior is a mixture of two Laplace distributions with scale parameters 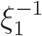 and 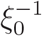, respectively, and *κ*_*mm*_′ a 0-1 indicator of whether precision element (*m, m*^′^) is substantially large. Through the use of separate hyperparameters for the active and null set, the mSSL avoids overshrinkage issues for elements in the active set commonly encountered in optimization-based strategies. As is typical for spike-slab frameworks, Beta-Bernoulli priors are placed on elements of *η* and *δ*:

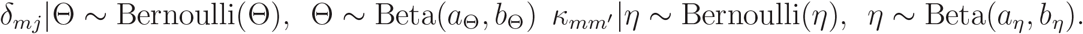

### Model fitting

The mSSL uses an Expectation Conditional Maximization algorithm to produce a sequential series of optimized point estimates for both coefficients ***β*** and inter-edge precision matrix Ω, where each solution was “warm-started” at the previous solution. This optimization strategy is equivalent to maximizing a posterior mode for a given specification of hyperparameters (*a*_*θ*_, *b*_*θ*_, *a*_*η*_, *b*_*η*_, *λ*_1_, *λ*_0_, *ξ*_1_, *ξ*_0_). The algorithm first holds (Ω, ***η***) constant and updates covariate coefficient matrix (***β, η***) via refined coordinate ascent and a Newton algorithm. Then, given these estimates, the algorithm updates ***η*** (closed form) and inter-edge dependence matrix Ω, which reduces to solving a graphical lasso problem (Friedman et al., 2008). The sequence is defined via ladders for *λ*_0_ and 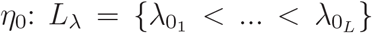 and [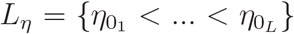} and a final solution is obtained when estimates of ***β*** and Ω are determined to have empirically converged (Deshpande et al., 2019).

### Inference

ConnReg’s inferential aims are to rigorously select which covariates are associated with changes in functional connectivity networks and to characterize which particular network edges drive significant covariate effects on functional connectivity. To select covariates, we propose a permutation test that produces multiplicity adjusted p-values for which covariates are significantly associated with functional connectivity. Our method permutes rows of our predictor matrix *X* to generate a null distribution 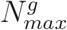 for a given regression method and then computes a test statistic *N*_*j*_ on unperturbed data. To determine which network edges drive these covariate effects, we produce stability selection scores with a computationally tractable bootstrap-based procedure that gives network edges a natural ordering in terms of importance when characterizing a covariate effect. As a secondary benefit, our stability selection procedure can be applied to precision matrix Ω to create a measure of confidence for which edges display inter-edge within network dependence. Our **Algorithms**, seen at the end of this manuscript with other figures and tables, fully outline our permutation test and stability selection procedure.

### Assessment of results in space of correlation matrices

To see the effects of covariates in the space of correlation matrices, we compare differences in predicted connectivity networks at distinct values of a covariate, holding all other covariates constant: ***R***_Δ_ = *g*^−1^{***µ***(***X***_*i*_ = ***x***_2_)} − ***g***^−1^{***µ***(***X***_*i*_ = ***x***_1_)}. This assessment of covariate-driven graph differences between networks is an active research area with many applications in neuroscience, as reviewed in recent work by Tomlinson et al. (2022). For example, one could compare differences in predicted graphs for a dichotomous case-control covariate while holding other covariates at their mean values. For our approach, we will see that the graph differences observed in the Fisher space comprise the predominate effects observed in the original space.

## 3. Simulation Studies

We conduct simulation studies to investigate the performance of our proposed methods. To reflect uncertainty in the estimation of subject-specific networks, we estimate each of *n* ‘observed’ subject-specific correlation matrices from 1000 draws of a multivariate normal distribution MVN(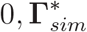), where 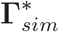 is a known *q* × *q* subject-specific correlation matrix constructed to vary by covariates. Γ_*sim*_ is generated in the Fisher space for all subjects, where Gaussian assumptions are justified, correlation matrix constraints are automatically satisfied, and inter-edge dependence among 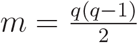 network edges can be fully controlled. The Fisher-space generative model used to produce *n* × *m* dataset Γ_*sim*_ is given by Γ_*sim*_ = ***Xβ***+***E***. Each row of Γ_*sim*_ induces a unique *q*×*q* true subject-specific correlation matrix 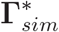. Subject-specific residuals ***E*** are generated by *MV N* (0_*M*_, Λ_*ρ*_), where Λ_*ρ*_ directly controls the level of inter-edge dependence in the dataset. We assume Λ_*ρ*_ follows a Toeplitz structure: 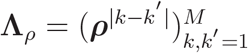. Covariates X are produced from *n* draws of *MV N* (0_*p*_, ***I***_*p*_).

In this simulation, we fix the number of edges to *m* = 45 (corresponding to a 10 × 10 correlation network) and the number of covariates to *p* = 30 while varying sample size (*n* = 150, 250) and level of inter-edge dependence (*ρ* = 0.9, 0.7, 0.5, 0.2). For each setting of *n* and *ρ* considered, we simulate 100 datasets. All results for *n* = 150 are included in Web Appendix A.1 as well as a setting with inter-edge dependence structure resembling our real data (i.e. non-Toeplitz structure including positive and negative second-order correlations). For every simulated dataset, a sparse ***β*** is generated as follows. First, 30 % of edges are randomly selected to have any signal with covariates. Next, for each selected edge 30 % of covariates are randomly selected to have non-zero signal with that edge. 5 covariates are then randomly selected to have 0 effect on any edges. For selected elements of ***β***, 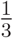 are randomly drawn from runif[0.1, 0.5], 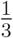 are randomly drawn from runif[0.5, 1], and ^1^ are randomly drawn from runif[1, 1.5] to assess our method’s robustness to capturing signals of varying strength. We next assess our method’s performance on our inferential framework. To determine the impact of accounting for second-order dependence, we compare our recommended approach against two modified versions of our framework that run single response sparse regressions on each edge (using the well-known LASSO (Tibshirani, 1996) and Sorted L-One Penalized Estimation (SLOPE) (Bogdan et al., 2015)) that do not take the presence of inter-edge dependence into account. More information on the SLOPE’s optimization procedure is available in Web Appendix A.1.

### Assessing inferential framework

**Table 1** illustrates the performance of our permutation-based hypothesis test and our bootstrap-based edge selection procedure. We compare our recommended approach (ConnReg_*mSSL*_) against two modified versions of our framework (ConnReg_*SLOPE*_, ConnReg_*LASSO*_) that do not take second order dependence into account. For our permutation-based hypothesis test, we assess power and experiment-wise type 1 (*T*_1_) error. To assess edge-level selection, 200 bootstrap samples were run for each simulated dataset considered. We compared partial area under the receiver operating characteristic curve (pAUC) and true positive rate when false positive rate = 0.01. We also estimate the stability selection cutoff needed to achieve a false positive rate = 0.01 for each method. Finally, we compare estimation accuracy for regression coefficient ***β*** across all three methods. ConnReg_*mSSL*_ demonstrates higher power and lower experiment-wise *T*_1_ error at *ρ* = 0.2, *ρ* = 0.5, *ρ* = 0.7 and *ρ* = 0.9 (with differences becoming more stark as *ρ* increases).

**Table 1.** Simulation results for sample size n = 250 assessing the operating characteristics of our framework in terms of hypothesis testing, edge-level selection, and estimation accuracy for one joint and two edge-specific regression approaches. Results are computed over 100 simulated datasets. Power and experiment-wise type 1 (T_1_) error are reported for hypothesis tests while pAUC at false positive rate = 0.01, the true positive rate at false positive rate = 0.01, and the stability threshold needed to attain false positive rate = 0.01 are reported for edge level selection. For estimation accuracy, the mean squared error of β relative to the mean squared error of OLS is reported.

| $\rho$ | method | Hypothesis Testing | | Edge Selection | | | Estimation |
| --- | --- | --- | --- | --- | --- | --- | --- |
| | | power | $T_1$ error | pAUC<sub>01</sub> (se) | TP<sub>01</sub> (se) | thresh<sub>01</sub> (se) | $\beta_{MSE}$ (se) |
| 0.9 | ConnReg<sub>mSSL</sub> | **0.997** | **<0.004** | **0.972** (0.015) | **0.987** (0.013) | 0.034 (0.010) | **0.117** (0.037) |
|  | ConnReg<sub>SLOPE</sub> | 0.762 | 0.04 | 0.871 (0.026) | 0.928 (0.024) | 0.364 (0.063) | 1.179 (0.151) |
|  | ConnReg<sub>LASSO</sub> | 0.05 | 0.10 | 0.849 (0.056) | 0.945 (0.020) | 0.945 (0.021) | 0.350 (0.048) |
| 0.7 | ConnReg<sub>mSSL</sub> | **0.997** | **0.01** | **0.947** (0.017) | **0.968** (0.016) | 0.059 (0.012) | **0.155** (0.046) |
|  | ConnReg<sub>SLOPE</sub> | 0.87 | 0.04 | 0.876 (0.025) | 0.933 (0.023) | 0.359 (0.043) | 1.134 (0.118) |
|  | ConnReg<sub>LASSO</sub> | 0.36 | 0.49 | 0.849 (0.050) | 0.942 (0.022) | 0.948 (0.014) | 0.327 (0.043) |
| 0.5 | ConnReg<sub>mSSL</sub> | **0.992** | **0.02** | **0.925** (0.018) | **0.948** (0.018) | 0.064 (0.012) | **0.195** (0.044) |
|  | ConnReg<sub>SLOPE</sub> | 0.935 | 0.04 | 0.874 (0.021) | 0.928 (0.021) | 0.363 (0.041) | 1.131 (0.113) |
|  | ConnReg<sub>LASSO</sub> | 0.545 | 0.61 | 0.847 (0.047) | 0.937 (0.021) | 0.947 (0.014) | 0.331 (0.043) |
| 0.2 | ConnReg<sub>mSSL</sub> | **0.990** | **<0.004** | **0.908** (0.023) | 0.935 (0.024) | 0.067 (0.011) | **0.246** (0.044) |
|  | ConnReg<sub>SLOPE</sub> | 0.939 | 0.06 | 0.875 (0.025) | 0.933 (0.023) | 0.362 (0.036) | 1.142 (0.112) |
|  | ConnReg<sub>LASSO</sub> | 0.674 | 0.69 | 0.848 (0.051) | **0.942** (0.021) | 0.949 (0.014) | 0.329 (0.040) |

ConnReg_*SLOPE*_ also produces competitive results at *ρ* = 0.2 and *ρ* = 0.5, suggesting its utility in settings where our framework is applied to data with low inter-edge dependence or with high dimension where computational speed is important. For edge selection, ConnReg_*mSSL*_ displayed superior true positive rates at the edge level while maintaining a low false positive rate of 0.01 at *ρ* = 0.5, 0.7, and 0.9. ConnReg_*mSSL*_ also demonstrated superior pAUC for all settings of *ρ* considered. Finally, each method also demonstrated different stability selection thresholds needed to obtain a false positive rate of 0.01; ConnReg_*mSSL*_ only required stability thresholds under 0.1 for all values of *ρ* while comparatively the ConnReg_*LASSO*_ required thresholds over 0.9, suggesting that when run without stability selection ConnReg_*LASSO*_ tends to be liberal and ConnReg_*mSSL*_ conservative in edge selection.

In terms of estimation accuracy, ConnReg_*mSSL*_ showed superior average *β*_*MSE*_ at all levels of *ρ* considered. Estimation accuracy of ConnReg_*SLOPE*_ and ConnReg_*LASSO*_ can be improved by using a two-step procedure where selection is first performed followed by OLS to reduce bias - ConnReg_*mSSL*_ does not have an ad-hoc two-step alternative. Estimation accuracy results using these two-step procedures are shown in Web Appendix A.1 and find that ConnReg_*LASSO*_ and ConnReg_*SLOPE*_ perform competitively at low levels of *ρ*. For ConnReg_*LASSO*_ *λ* was selected by 10-fold cross-validation minimizing MSE.

In summary, simulation results provide evidence for the validity of our inferential procedure. For our designed permutation hypothesis test, in the presence of inter-edge dependence ConnReg_*mSSL*_ dominates in terms of power and experiment-wise *T*_1_ error compared to approaches running independent edge-specific regressions. Similarly, for our bootstrap-based edge selection procedure in the presence of inter-edge dependence our recommended approach shows superior true positive rates and higher pAUC at fixed false positive rates. Finally, the superior estimation accuracy of ***β*** by ConnReg_*mSSL*_ in the presence of inter-edge dependence demonstrates the tangible benefits of accounting for second order associations when regressing subject-specific networks on covariates. Our framework also performed well with faster parallel-edge specific regression approaches when inter-edge dependence was low, suggesting it can effectively be used to perform rigorous inference for applications with low inter-edge dependence or for applications where the practitioner wants to scale up to networks with very high dimensions. We note that results from all simulations conducted with sample size *n* = 150 and with dependence structure matching our Human Connecctome Project data set (see Web Appendix A.1) demonstrate the same beneficial properties for our approach as those observed in the main body of this manuscript for *n* = 250.

## 4. Analyzing the HCP Young Adult Study

We apply ConnReg to characterize heterogeneity in connectivity patterns of 1003 subjects taken from the Human Connectome Project (HCP) Young Adult Study (Van Essen et al., 2013). Subjects are healthy individuals between the ages of 22-35 years old with fMRI BOLD score time series summarized for 15 aggregated functional brain regions (Glasser et al., 2016; Robinson et al., 2018). Detailed information on data retrieval, pre-processing steps, and a comprehensive anatomical breakdown of each aggregated functional brain region is discussed in Web Appendices A.2.1 and A.2.2. We considered 7 covariates: two cognitive tests related to language processing ability, and the structural area of five brain regions related to cortical hubs connecting intrinsic functional connectivity sub-networks. Cortical hubs are hypothesized to be the means through which distinct sub-networks communicate with one another (Buckner et al., 2009; Liu et al., 2018). From the pre-processed time series, we form 15 × 15 subject specific networks and transform to the Fisher space to produce a 1003×105 stacked matrix of network edges. For hypothesis tests, 250 permutations were used to generate null distributions. An initial empirical assessment of our data (see Web Appendix A.2.3) indicated a high presence of inter-edge dependence within subject-specific networks. Similarly, an empirical comparison of within network inter-edge dependence in the Fisher and original space of correlation networks show highly similar patterns, in terms of both magnitude and direction. This high degree of similarity suggests that it may be reasonable to directly interpret second-order associations found in the Fisher space.

### Linking phenotype with functional connectivity networks

The seven covariates considered are grouped into language processing assessments (“ReadEng”, “PicVocab”) and anatomical areas of intrinsic functional connectivity hubs (“Precuneus”, “Postcentral Gyrus”, “Cuneus”, “Posteriorcingulate”, “Temporalpole”). We find ReadEng, Precuneus area, and Postcentral Gyrus area (p-value: *<*0.004) to be significantly associated with changes in subject-specific functional connectivity networks and all other covariates to not be significantly associated (p-values: *>*0.996). Given covariates found to significantly vary with functional connectivity, we ran our bootstrap-based stability selection procedure to identify which edges were driving these effects. Figure 2 shows heatmaps demonstrating stability selection results over 200 bootstrap samples for covariate-network edge associations and second order edge-edge associations in the Fisher space. Stability selection results for second order associations validate the empirical examination performed at the beginning of this section, as many within-network edge-edge associations are found to be associated over 85 percent of the time (i.e. stability selection scores greater than 0.85). A comprehensive overview of first and second order stability selection results are shown in Web Appendix A.2.3.

**Figure 2.**
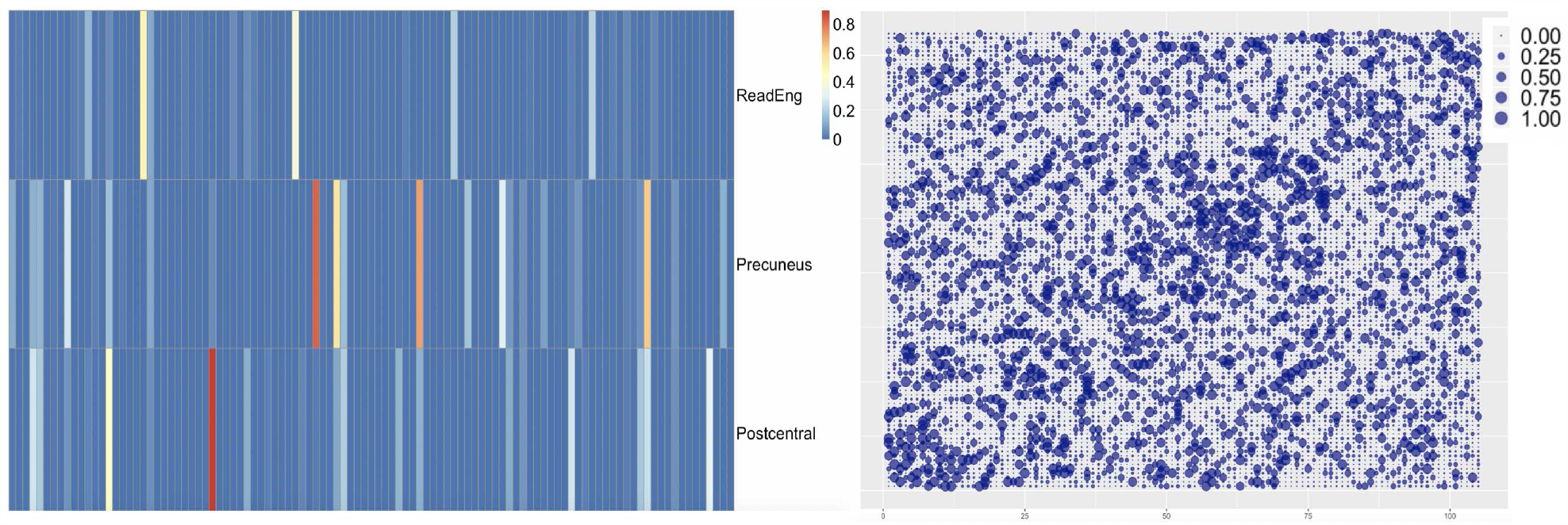
**Left Panel:** Bootstrap-based stability selection results for potential covariate-edge combinations over 200 bootstraps in Fisher space for covariates found to be significantly associated with functional connectivity networks. The heatmap represents the proportion of times a covariate-edge combination was selected across bootstrap samples. **Right Panel:** Over the same 200 bootstrap samples, bootstrap-based stability selection results for all potential second-order edge-edge associations in Fisher space.

Holding all other covariates at mean levels, to characterize a covariate’s impact on functional connectivity we assess graph differences for predicted networks at the *X*_0.1_ percentile and the *X*_0.9_ percentile in both the Fisher space and the space of correlation networks. Figure 3 displays graph differences for our three flagged covariates. Results are overlaid with stability selection proportions conducted for each covariate-edge combination in Fisher space. We observe that salient features are preserved in both Fisher space and correlation space with respect to direction and magnitude of edge differences. Predicted network differences are sparse in the Fisher space by construction, and approximately sparse in the original correlation space but with some near zero magnitude edges appearing when transformed from the Fisher space. The results provide strong evidence that the primary edges driving functional connectivity changes can be identified in Fisher space both in terms of magnitude and direction (increasing or decreasing) - the dominant edge differences identified in Fisher space in Figure 3 are the same sign (color), similar magnitude (thickness), and the same direction (solid vs. dashed) when transformed to the original space of correlation networks for each significant covariate.

**Figure 3.**
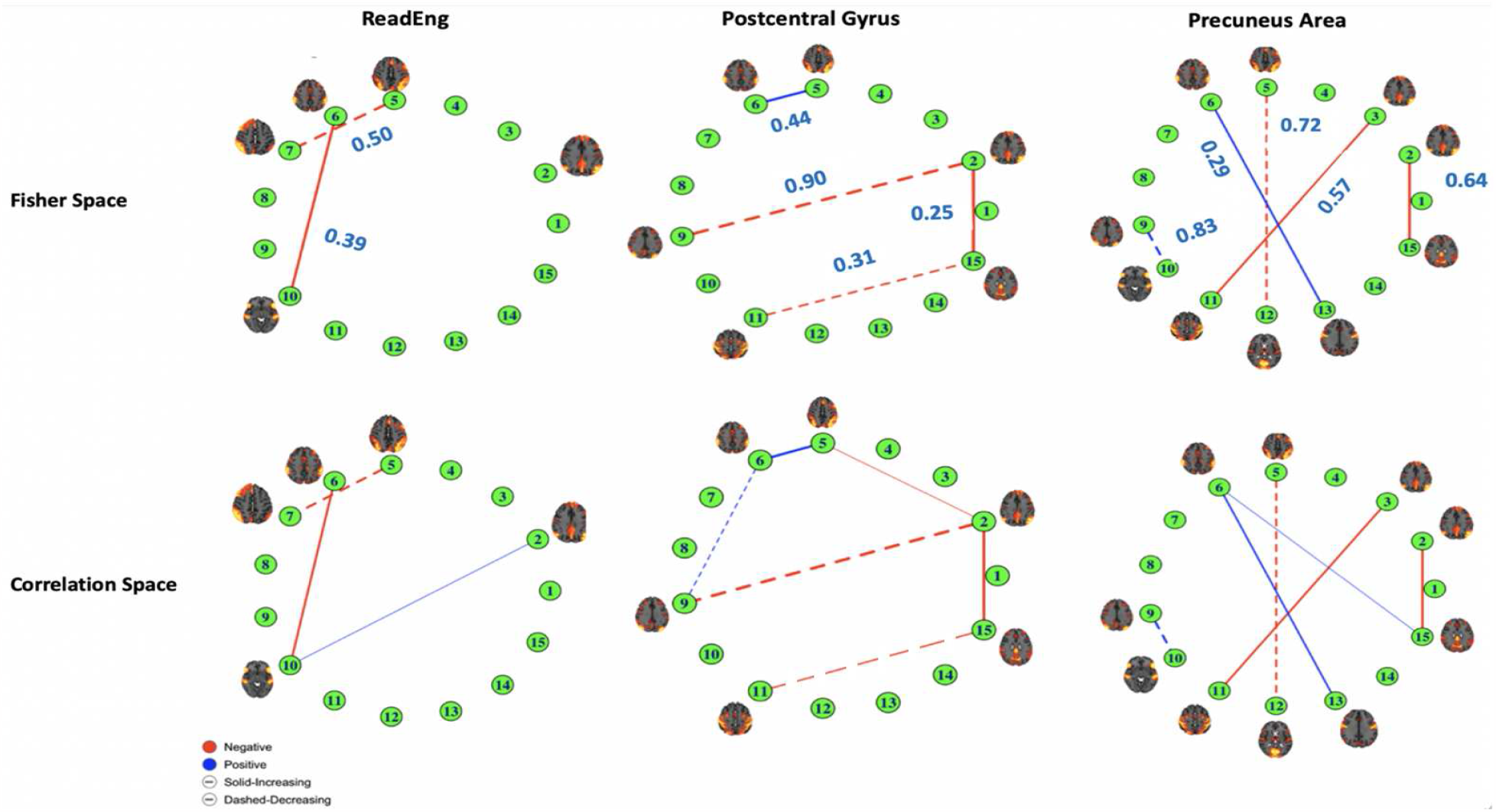
**Top Row:** The difference between predicted networks ***R***_Δ_ = *g*^−1^{***µ***(***X***_*i*_ = ***x***_0.9_)} − ***g***^−1^{***µ***(***X***_*i*_ = ***x***_0.1_)} in Fisher space for flagged covariates ReadEng, Postcentral Gyrus Area, and Precuneus Area. Edge widths represent the magnitude of edge difference between predicted networks. Edge color indicates whether the difference between a predicted edge at the 90th and 10th percentile was negative or positive. Finally edge type (solid or dashed) indicates whether the predicted edge at the 90th percentile was smaller or greater in magnitude than the corresponding predicted edge at the 10th percentile. Flagged edges are ovarlaid with covariate-edge stability selection scores over 200 bootstrap samples. **Bottom Row:** The difference between predicted networks in the space of correlation networks for covariates ReadEng, Postcentral Gyrus Area, and Precuneus Area. Relevant nodes are supplemented with images corresponding to their anatomical areas.

For ReadEng, prominent edges involve nodes that correspond to the cerebellum, amygdala, brain stem, thalamus, and putamen. Combined with known functions of these subcortical structures, recent research implicating the cerebellum’s role in language recognition indicates that predicted graph differences may link language recognition both with emotional response and with episodic memory and emotional expression (Koziol et al., 2014; Basinger and Hogg, 2019; Torrico and Munakomi, 2019). For Precuneus Area, the most prominently flagged edges involve nodes that correspond to the hippocampus, cerebellum, amygdala, thalamus, diencephalon, and brain stem. Given the function of the precuneus, edges seem to link episodic memory and motor movement, addiction, and emotion (Fauchon et al., 2019; Tomasi and Volkow, 2020; Berkovich-Ohana et al., 2020). The most prominent edge flagged for Postcentral Gyrus Area involves nodes that correspond to the brain stem, cerebellum, and hippocampus. Given the function of the postcentral gyrus, the majority of edges seem to link memory with motor movement, with the association between node 5 and node 6 possibly linking emotional response with episodic memory (DiGuiseppi and Tadi, 2020; Fu et al., 2019). A more comprehensive biological interpretation of prominent flagged edges for all significant covariates is given in Web Appendix A.2.3.

### Interpretation of second-order dependence

Given the presence of substantial inter-edge dependence, Figure 4 illustrates the mostly highly correlated edges (those with stability selection = 1 and partial correlation ⩾ 0.7). The cerebellum, heavily represented in node 8, has a central role in regulating associations between anatomical areas and appears to act as a “hub” of strong second order associations. A comprehensive overview of all potential second order associations, in terms of stability selection and partial correlation, is provided in Web Appendix A.2.3 along with a more comprehensive interpretation of the second order associations highlighted in Figure 4.

**Figure 4.**
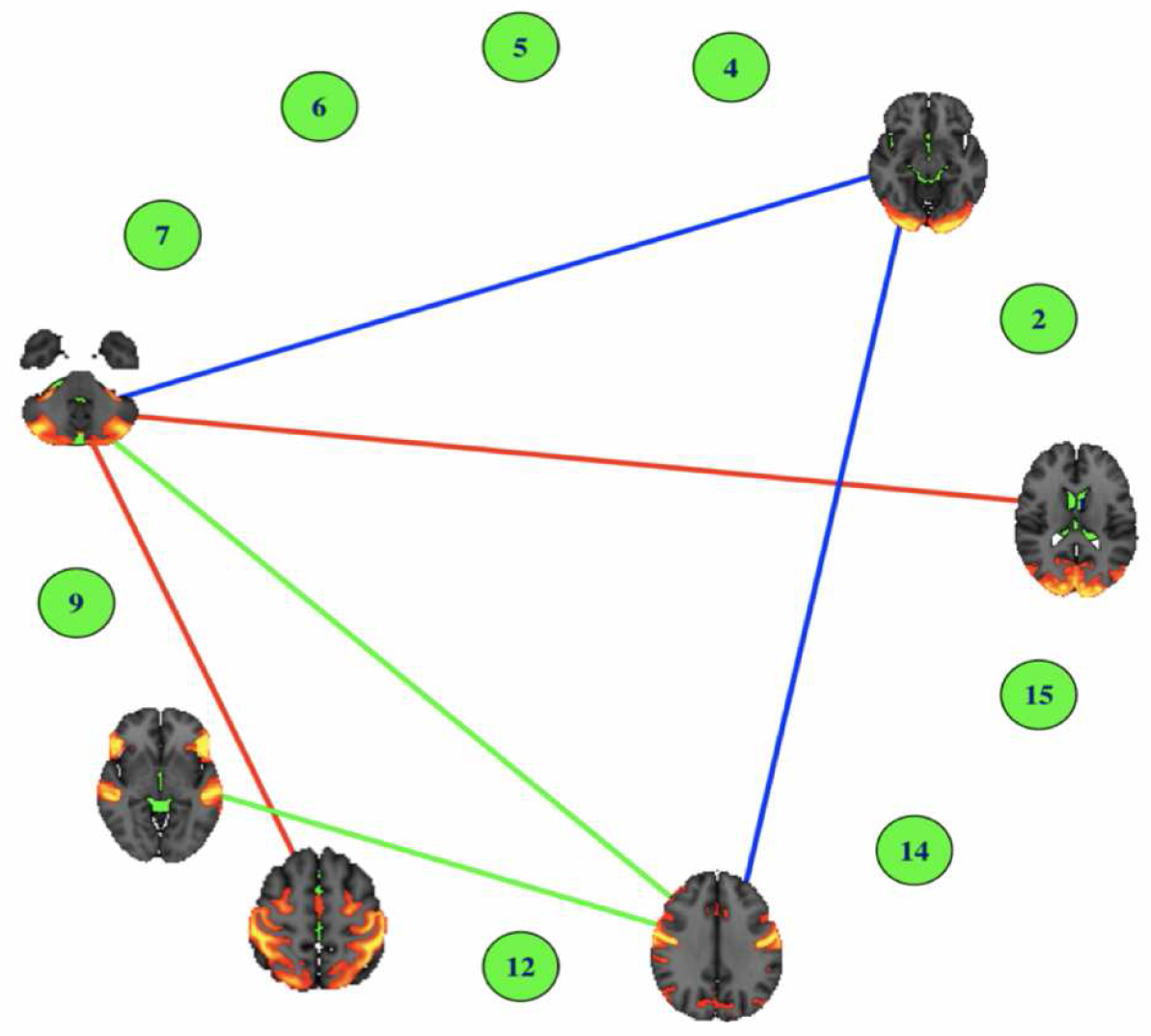
The top second order associations (stability selection = 1 and partial correlation ⩾ 0.7) are shown. Edge colors indicate correlated edges and the image of functional brain regions corresponding to selected edges are overlaid on top of the appropriate network nodes.

## 5. Discussion

In this article we introduce a principled framework for regressing entire correlation networks on covariates, connectivity regression (ConnReg), motivated by applications in functional connectivity. We utilize a sparse multivariate linear model in a transformed Fisher space that enables unconstrained modeling on the real line, justifiable Gaussian assumptions, and automatically satisfied correlation matrix constraints. Via the precision matrix in the transformed Fisher space, our framework is able to fully estimate and account for within network inter-edge dependence, which is shown via simulations to lead to gains in estimation efficiency and selection accuracy. We introduced a permutation hypothesis test that produces multiplicity-adjusted p-values for which covariates vary with functional connectivity networks and a bootstrap-based stability selection procedure that characterizes which network edges drive significant covariate effects. Results in the Fisher space are shown to be highly concordant with those transformed back to the space of correlation networks, aiding interpretability.

The fact that connectivity regression of a *p* × *p* network requires assessment of an 𝒪 (*p*^4^) second-order association matrix raises computational challenges to apply ConnReg_*mSSL*_ to very large networks. However, assessment of functional connectivity networks represented by lower dimensional embeddings is an active area of contemporary research (Casanova et al., 2021; Gallos et al., 2021; Gao et al., 2021), so networks of small to moderate size are clearly of interest. Also, ConnReg_*SLOPE*_ runs much faster, not requiring calculation of the 𝒪 (*p*^4^) second-order association matrix, and does not sacrifice too much performance relative to ConnReg_*mSSL*_ when the magnitude of second order dependence is not too high, so could be an alternative to consider for very high-dimensional networks. In future work, we will seek to improve the computational efficiency of our joint modeling algorithm to enable the benefits of accounting for second order associations while scaling to larger networks. Other future extensions can include the relaxation of linearity assumptions to allow connectivity to vary smoothly with continuous covariates while maintaining computational feasibility or the incorporation of random effects to adjust for inter-individual correlation structure when multi-level designs are encountered.

In conclusion, we view this manuscript’s framework and the idea of fully leverage within network inter-edge dependence as fundamental building blocks for new methods to be developed for the improved assessment of the link between functional connectivity and phenotypes.

## Supporting information

Web Appendix

## Supplementary Materials and Acknowledgements

Software is available at https://github.com/nmd1994/Connectivity-Regression. This work was partially supported by CA-178744 and CA-244845 from the National Cancer Institute, and TR-001878 from the National Center for Advancing Translational Science.

---

*Figure: Algorithms for Conducting Inference*

## Algorithm 1

Permutation Based Hypothesis Test

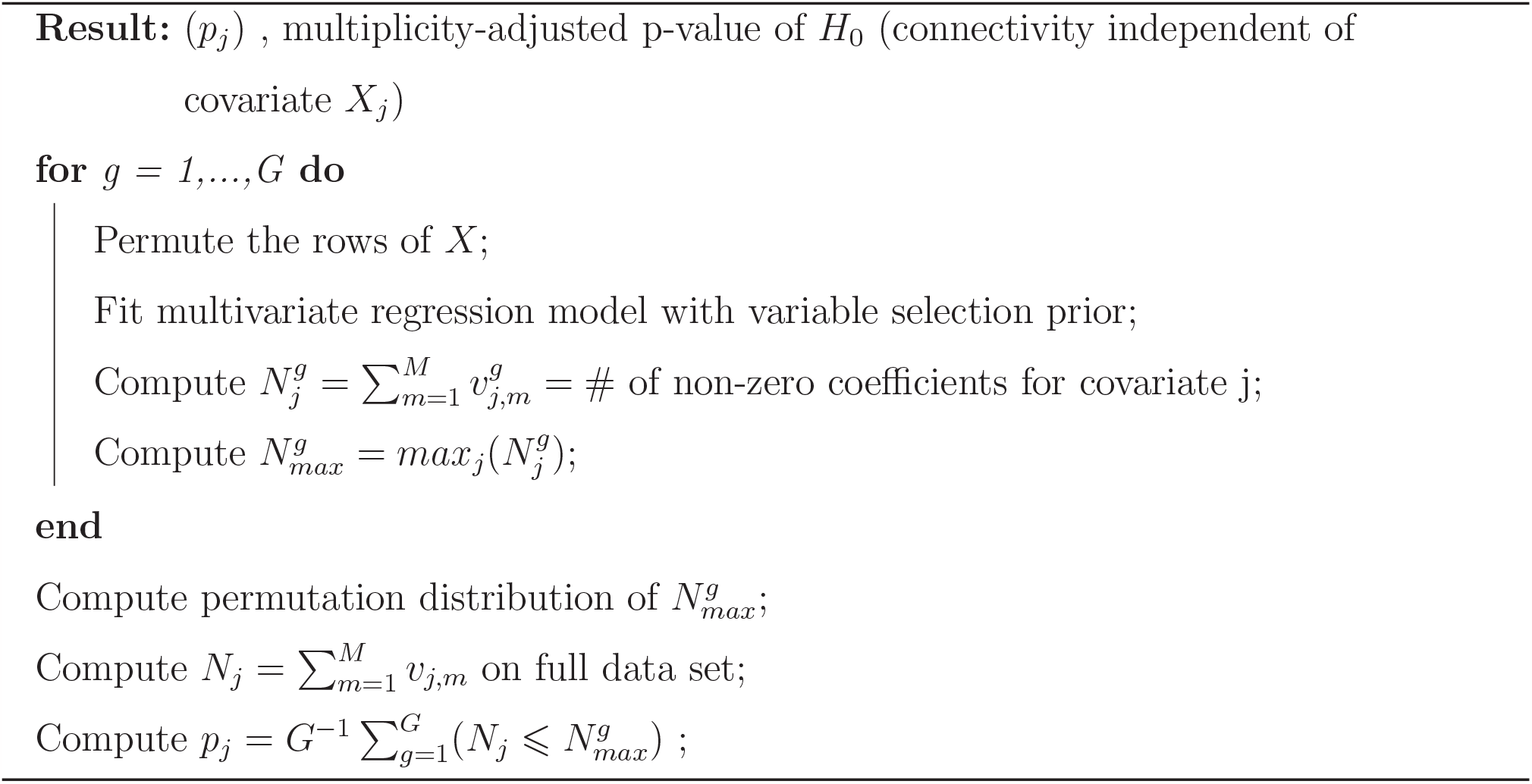

## Algorithm 2

Bootstrap-Based Stability Selection

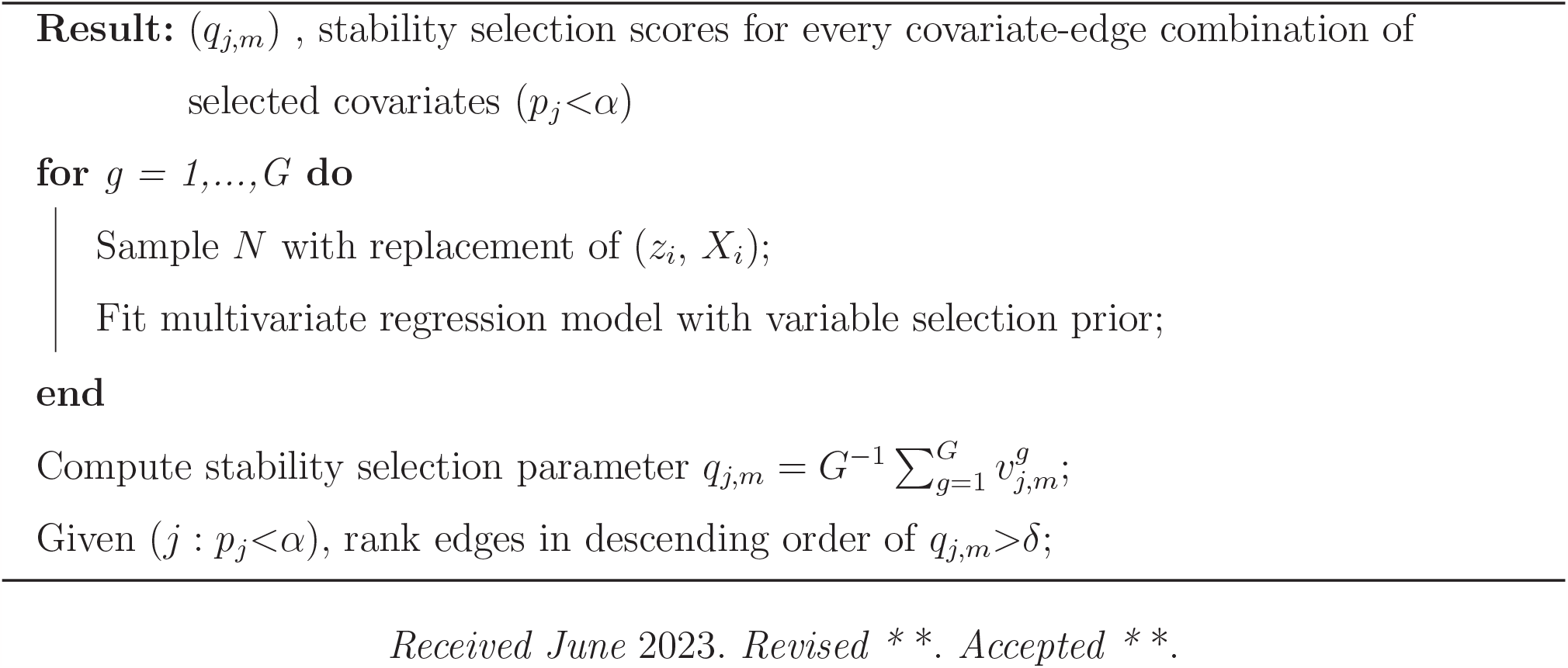

