## Supplementary material for "Connectivity Regression": Web Appendix

### A Web Appendix Materials

#### A.1 Additional Simulation Settings

**SLOPE formulation:** SLOPE is a convex problem whose solution via proximal gradient algorithms has similar computational complexity to the LASSO (Tibshirani 1996), and its optimization problem can be written as:

$$\min \ \frac{1}{2}||\gamma_m - X\beta_m||^2 + pen(\beta_m)$$

$$\text{where } \ pen(\beta_m) = \sum_{j=1}^{p} \lambda_j |\beta_m|_{(j)} \ \text{ and } \ \lambda_1 \geq \lambda_2 \geq ... \geq \lambda_p$$

$$\text{and } \ |\beta_m|_{(1)} \geq |\beta_m|_{(2)} \geq ... \geq |\beta_m|_{(p)} \ \text{ are decreasing absolute values of } \beta_{j,m}$$

In the case where $X$ is orthogonal, SLOPE is theoretically guaranteed to control false discovery rate with choice of lambda sequence $\lambda_{(BH)}(i) = z(1 - i \times \frac{q}{2p})$ with the Benjamin and Hochberg procedure (BH) designed to control false discovery rate. We propose the same permutation test and stability selection procedures described above to do inference for ConnReg$_{SLOPE}$ and ConnReg$_{LASSO}$.

**Simulations conducted for** $n = 250$ We first present the simulation results presented in the main body of this manuscript with one exception; the $\beta_{MSE}$ for the SLOPE and LASSO is now computed in two steps: selection is first performed followed by estimation via OLS. We note that the mSSL still exhibits superior $\beta_{MSE}$ at higher levels of second-order dependence $\rho$ while SLOPE and LASSO perform competitively in terms of estimation accuracy at lower levels of $\rho$.

| $\rho$ | method | Hypothesis Testing | | Edge Selection | | | Estimation |
|---|---|---|---|---|---|---|---|
| | | power | $T_1$ error | $\text{pAUC}_{01}$ (se) | $\text{TP}_{01}$ (se) | $\text{thresh}_{01}$ (se) | $\beta_{MSE}$ (se) |
| 0.9 | $\text{ConnReg}_{mSSL}$ | 0.997 | <0.004 | 0.972 (0.015) | 0.987 (0.013) | 0.034 (0.010) | 0.117 (0.037) |
| | $\text{ConnReg}_{SLOPE}$ | 0.762 | 0.04 | 0.871 (0.026) | 0.928 (0.024) | 0.364 (0.063) | 0.201* (0.039) |
| | $\text{ConnReg}_{LASSO}$ | 0.05 | 0.10 | 0.849 (0.056) | 0.945 (0.020) | 0.945 (0.021) | 0.214* (0.040) |
| 0.7 | $\text{ConnReg}_{mSSL}$ | 0.997 | 0.01 | 0.947 (0.017) | 0.968 (0.016) | 0.059 (0.012) | 0.155 (0.046) |
| | $\text{ConnReg}_{SLOPE}$ | 0.87 | 0.04 | 0.876 (0.025) | 0.933 (0.023) | 0.359 (0.043) | 0.188* (0.036) |
| | $\text{ConnReg}_{LASSO}$ | 0.36 | 0.49 | 0.849 (0.050) | 0.942 (0.022) | 0.948 (0.014) | 0.206* (0.035) |
| 0.5 | $\text{ConnReg}_{mSSL}$ | 0.992 | 0.02 | 0.925 (0.018) | 0.948 (0.018) | 0.064 (0.012) | 0.195 (0.044) |
| | $\text{ConnReg}_{SLOPE}$ | 0.935 | 0.04 | 0.874 (0.021) | 0.928 (0.021) | 0.363 (0.041) | 0.189* (0.032) |
| | $\text{ConnReg}_{LASSO}$ | 0.545 | 0.61 | 0.847 (0.047) | 0.937 (0.021) | 0.947 (0.014) | 0.205* (0.033) |
| 0.2 | $\text{ConnReg}_{mSSL}$ | 0.990 | <0.004 | 0.908 (0.023) | 0.935 (0.024) | 0.067 (0.011) | 0.246 (0.044) |
| | $\text{ConnReg}_{SLOPE}$ | 0.939 | 0.06 | 0.875 (0.025) | 0.933 (0.023) | 0.362 (0.036) | 0.190* (0.035) |
| | $\text{ConnReg}_{LASSO}$ | 0.674 | 0.69 | 0.848 (0.051) | 0.942 (0.021) | 0.949 (0.014) | 0.203* (0.033) |

Table 1: Simulation results for sample size $n = 250$ assessing the operating characteristics of our framework in terms of hypothesis testing, edge-level selection, and estimation accuracy for one joint and two edge-specific regression approaches. Results are computed over 100 simulated datasets. Power and experiment-wise type 1 ($T_1$) error are reported for hypothesis tests while pAUC at false positive rate = 0.01, the true positive rate at false positive rate = 0.01, and the stability threshold needed to attain false positive rate = 0.01 are reported for edge level selection. For estimation accuracy, the mean squared error of $\beta$ relative to the mean squared error of OLS is reported. For $\text{ConnReg}_{SLOPE}$ and $\text{ConnReg}_{LASSO}$, $\beta_{MSE}$ is computed in two steps: selection is first performed followed by estimation via OLS.

**Additional simulations conducted for $n = 150$:** Given the simulation results presented in the main body of this manuscript, we would like to assess our framework's performance for data of a different sample size. Given the high dimensional computations needed for a joint approach, we would like to see how our method performs when less subject-specific data is available. We assess, for the same simulation design described in section 3, performance for sample size $n = 150$. Similar to results obtained for $n = 250$, we see that ConnReg has superior performance in terms of power and $T_1$ error. Likewise, ConnReg has better operating characteristics for stability selection. For estimation accuracy, ConnReg has superior MSE for higher values of $\rho$. MSE for the SLOPE and LASSO are conducted in two-steps: selection is first performed followed by OLS.

| | | Hypothesis Testing | | Stability Selection | | | Estimation |
|---|---|---|---|---|---|---|---|
| $\rho$ | method | power | $T_1$ error | $\text{pAUC}_{01}$ (se) | $\text{TP}_{01}$ (se) | $\text{thresh}_{01}$ (se) | $\beta_{MSE}$ (se) |
| 0.9 | ConnReg | 0.995 | 0.02 | 0.909 (0.024) | 0.959 (0.018) | 0.026 (0.009) | 0.105 (0.029) |
| | SLOPE | 0.660 | 0.03 | 0.812 (0.034) | 0.873 (0.034) | 0.449 (0.073) | 0.198* (0.035) |
| | Lasso | 0.045 | 0.11 | 0.802 (0.062) | 0.942 (0.025) | 0.942 (0.025) | 0.206* (0.033) |
| 0.7 | ConnReg | 0.988 | <0.004 | 0.904 (0.023) | 0.933 (0.024) | 0.071 (0.015) | 0.153 (0.044) |
| | SLOPE | 0.805 | 0.01 | 0.806 (0.032) | 0.868 (0.033) | 0.447 (0.056) | 0.194* (0.035) |
| | Lasso | 0.353 | 0.480 | 0.791 (0.054) | 0.887 (0.028) | 0.945 (0.017) | 0.199* (0.035) |
| 0.5 | ConnReg | 0.988 | <0.004 | 0.875 (0.025) | 0.908 (0.026) | 0.086 (0.016) | 0.197 (0.039) |
| | SLOPE | 0.923 | 0.07 | 0.874 (0.021) | 0.870 (0.027) | 0.442 (0.041) | 0.191* (0.029) |
| | Lasso | 0.543 | 0.62 | 0.804 (0.045) | 0.891 (0.028) | 0.946 (0.013) | 0.194* (0.030) |
| 0.2 | ConnReg | 0.982 | <0.004 | 0.847 (0.030) | 0.883 (0.028) | 0.083 (0.014) | 0.253 (0.044) |
| | SLOPE | 0.919 | 0.130 | 0.808 (0.031) | 0.870 (0.031) | 0.445 (0.044) | 0.187* (0.031) |
| | Lasso | 0.637 | 0.730 | 0.804 (0.049) | 0.892 (0.031) | 0.945 (0.012) | 0.194* (0.027) |

Table 2: Simulation results for sample size $n = 150$ assessing the operating characteristics of our framework in terms of hypothesis testing, edge-level selection, and estimation accuracy for one joint and two edge-specific regression approaches. Results are computed over 100 simulated datasets. Power and $T_1$ error are reported for hypothesis tests while pAUC at false positive rate = 0.01, the true positive rate at false positive rate = 0.01, and the stability threshold needed to attain false positive rate = 0.01 are reported for edge level selection. For estimation accuracy, the mean squared error of $\beta$ relative to the mean squared error of OLS is reported. For the SLOPE and LASSO, $\beta_{MSE}$ is computed in two steps: selection is first performed followed by estimation via OLS.

To assess our inferential framework, we also consider an additional simulation setting for sample size $n = 150$ generated from a subset of our real data ($q = 45$ edges). In this setting, second order dependence (within-network dependence among edges in the Fisher space) is set to be a subset of the empirical second order correlation from our dataset in the Human Connectome Project (see Web Appendix A.2 for the real data analysis section). To generate realistic covariate associations, we set the sparsity patterns (as well as signs and magnitudes) to those obtained from running the SLOPE independently on the brain regions considered, and generate 100 random datasets in the Fisher space in the same manner as our previous simulations. Covariates are set to $p = 7$ as in our real data analysis. This setting is designed to demonstrate the robustness of Connectivity Regression to less structured settings with both negative and positive second order associations. We note that our approach (ConnReg) reports notably superior power and pAUC versus the approaches that don't take second order dependence into account while maintaining excellent $T_1$ error control.

| | Hypothesis Testing | | Stability Selection | |
|---|---|---|---|---|
| method | power | $T_1$ error | $\text{pAUC}_{01}$ (se) | $TP_{01}$ (se) |
| ConnReg | 1 | <0.004 | 0.963 (0.015) | 0.991 (0.011) |
| Lasso | 0.324 | 0.180 | 0.763 (0.104) | 0.986 (0.069) |
| SLOPE | 0.863 | <0.004 | 0.799 (0.042) | 0.989 (0.015) |

Table 3: Hypothesis testing and stability selection conducted for $n = 150$ with second order dependence structure derived from the second order dependence structure set to the empirical correlation from the Human Connectome Project. Results are computed over 100 simulated datasets. Power and T1 error are reported for hypothesis tests while pAUC at false positive rate = 0.01, the true positive rate at false positive rate = 0.01, and the stability threshold needed to attain false positive rate = 0.01 are reported for edge level selection.

We also explore, at sample size $n = 150$, how *ConnReg* performs in terms of estimation accuracy when the mSSL is replaced by ad-hoc multivariate approaches. We use the simulated datasets in Table 2 along with a setting where $q = 45$ edges have a 3 block-diagnoal dependence structure with $\rho = 0.7$, $\rho = 0.5$, and $\rho = 0.2$. Ad-hoc approaches ("SLOPE-OLS-SUR, Lasso-OLS-SUR"), were fit in the following procedure: $q$ single response selections were performed and refitted with OLS, an empirical covariance matrix was computed from the residuals, and the subset of selected responses were refit with generalized least squares applied to a system of equation (i.e. seemingly unrelated regressions (SUR)). We note that using the mSSL still performs better for high levels of second order dependence ($\rho = 0.9, \rho = 0.7$) while the ad-hoc approaches do well at lower levels of residual dependence. We caution that using 'ad-hoc' approaches do not gain any of the inferential benefits demonstrated by the mSSL for selection of covariates or edges.

| | | Estimation |
|---|---|---|
| $\rho$ | method | $\beta_{MSE}$ |
| 0.9 | mSSL | 0.106 (0.030) |
| | SLOPE-OLS-SUR | 0.156 (0.036) |
| | Lasso-OLS-SUR | 0.129 (0.029) |
| 0.7 | mSSL | 0.154 (0.045) |
| | SLOPE-OLS-SUR | 0.171 (0.037) |
| | Lasso-OLS-SUR | 0.155 (0.032) |
| 0.5 | mSSL | 0.196 (0.038) |
| | SLOPE-OLS-SUR | 0.180 (0.029) |
| | Lasso-OLS-SUR | 0.173 (0.028) |
| 0.2 | mSSL | 0.252 (0.044) |
| | SLOPE-OLS-SUR | 0.188 (0.032) |
| | Lasso-OLS-SUR | 0.192 (0.028) |
| block | mSSL | 0.157 (0.041) |
| | SLOPE-OLS-SUR | 0.172 (0.039) |
| | Lasso-OLS-SUR | 0.153 (0.032) |

Table 4: Esimation accuracy results for using the mSSL or ah-hoc alternatives over 100 simulated datasets at $n = 150$. Inter-edge dependence is set at $\rho = 0.9, 0.7, 0.5, 0.2$ and a block-diagonal with blocks at $\rho = 0.7, 0.5, 0.2$. $\beta_{MSE}$ for each approach is reported as a ratio relative to OLS.

Simulation results assessing the operating characteristics of our framework show the same patterns for all sample sizes ($n = 250, n = 150$) considered. The superior performance for the mSSL at higher values of $\rho$ demonstrate the benefits of accounting for inter-edge dependence within subject-specific networks. Likewise, the competitive performance of the SLOPE at lower levels of $\rho$ demonstrate how our framework can be adapted for use in applications where inter-edge dependence is low or in settings where subject-specific networks are high dimensional.

We next provide supporting information for our analysis of 1003 subject-specific functional connectivity networks taken from the Human Connectome Project. Section A.2.1 describes how the data was obtained and subject specific networks were formed. Section A.2.2 provides a breakdown of the anatomical makeup for each of the 15 "nodes" of our subject-specific networks. Finally, sections A.2.3 provides more comprehensive supporting information for the results in section four in the main body of the manuscript.

#### A.2 Human Connectome Project

##### A.2.1 Instructions for Estimating Subject-Specific Correlation Matrices and Extracting Anatomical Details for ICA Components

With the creation of a free ConnectomeDB account, timeseries with 4800 timepoints for each of 1003 subjects was obtained from the "HCP1200 Parcellation + Timeseries + Netmaps" data bundle ("https://www.humanconnectome.org/study/hcp-young-adult/docu ment/extensively-processed-fmri-data-documentation") - analysis for this paper was performed with the data divided into 15 independent components included in this bundle. Independent components, which each serve as aggregated brain regions, are determined through extensive pre-processing via a combination of group-PCA and group-ICA steps. Given 4800 timepoints, empirical $15 \times 15$ subject-specific correlation matrices were estimated for each of 1003 subjects. Information for the anatomical makeup of each IC (shown next) can be extracted from the Grayordinates data in the bundle described above. Documentation for the files in this bundle can be obtained here: ("https://www.humanconnectome.org/storage/app/media/documentation/s1200/HCP1200-DenseConnectome+PTN+Appendix-July2017.pdf").

##### A.2.2 Anatomical Makeup of ICA nodes

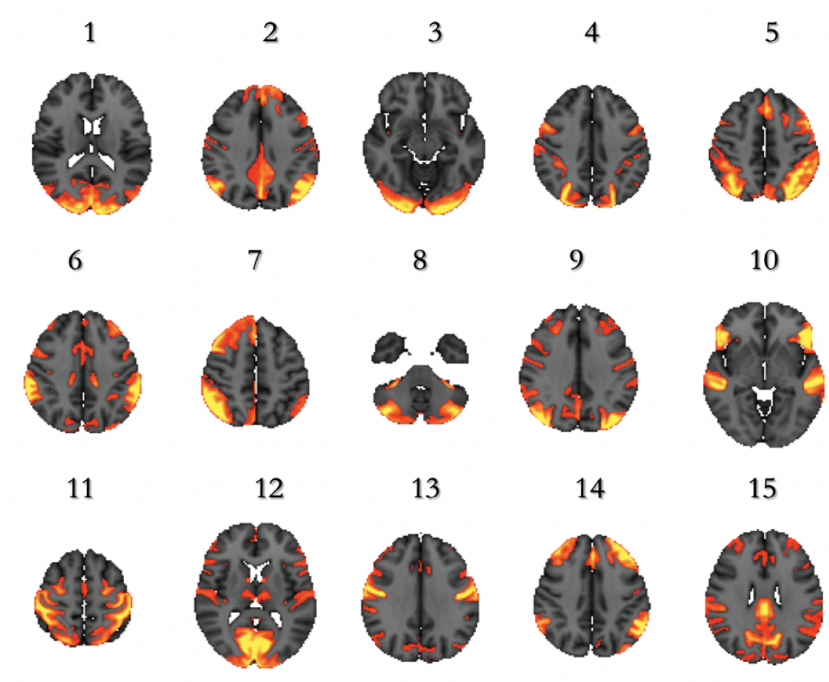


Figure 1: Highlighted regions on axial slices of the brain indicate which anatomical regions correspond to each ICA node in our analysis.

Given the images corresponding to each ICA node, we next show the percentage makeup of each ICA node in terms of particular anatomical brain structures. Information is obtained from the files ”melodic-IC.dscalar.nii” and ”melodic-IC-ftb.dlabel.nii” using the Rpackage ”ciftiTools”. Table 3 summarizes the anatomical breakdown of each node below.

| IC (node) | Accumbens | Amygdala | Brain Stem | Caudate | Cerebellum | Diencephalon | Hippocampus | Pallidum | Putamen | Thalamus |
|---|---|---|---|---|---|---|---|---|---|---|
| 1 | 0 | 0.439 | 0.16 | 0 | 0.019 | 0 | 0.32 | 0 | 0 | 0.063 |
| 2 | 0 | 0.002 | 0.193 | 0.01 | 0.567 | 0.038 | 0.144 | 0 | 0 | 0.045 |
| 3 | 0.158 | 0 | 0.392 | 0 | 0 | 0.129 | 0 | 0.321 | 0 | 0 |
| 4 | 0.147 | 0 | 0.084 | 0.016 | 0.299 | 0.012 | 0.004 | 0.024 | 0.331 | 0.084 |
| 5 | 0 | 0.073 | 0.049 | 0.044 | 0.692 | 0.005 | 0.063 | 0.027 | 0.019 | 0.028 |
| 6 | 0 | 0.097 | 0.138 | 0 | 0.086 | 0.014 | 0 | 0.128 | 0.241 | 0.297 |
| 7 | 0.002 | 0.001 | 0.251 | 0.024 | 0.584 | 0.038 | 0.03 | 0.015 | 0.015 | 0.041 |
| 8 | 0.1 | 0.002 | 0.095 | 0.046 | 0.7 | 0.049 | 0.008 | 0.022 | 0.012 | 0.056 |
| 9 | 0.01 | 0.038 | 0.055 | 0 | 0.309 | 0.016 | 0.557 | 0 | 0 | 0.016 |
| 10 | 0 | 0.219 | 0.041 | 0.02 | 0.653 | 0.009 | 0.014 | 0.018 | 0.001 | 0.024 |
| 11 | 0 | 0 | 0.064 | 0 | 0.782 | 0.007 | 0.05 | 0 | 0.004 | 0.093 |
| 12 | 0 | 0 | 0.182 | 0.048 | 0.135 | 0.223 | 0.001 | 0.015 | 0.004 | 0.391 |
| 13 | 0.002 | 0 | 0.018 | 0.038 | 0.177 | 0.049 | 0.025 | 0.015 | 0.546 | 0.071 |
| 14 | 0.0149 | 0 | 0.076 | 0.2 | 0.594 | 0.019 | 0.002 | 0.02 | 0.045 | 0.029 |
| 15 | 0.127 | 0 | 0.407 | 0.002 | 0.052 | 0.052 | 0.251 | 0.02 | 0.045 | 0.043 |

Table 5: Breakdown of each ICA node in terms of anatomical structures. The number between 0-1 represents the proportion of that node that is represented by a particular anatomical structure.

##### A.2.3 Supporting information for real data analysis conducted from the Human Connectome Project (HCP)

We begin this section by providing the empirical second-order correlation matrix discussed in the beginning of section 4 of this manuscript. We note that the empirical associations in the Fisher space and the original space of correlation networks show highly similar patterns, both in terms of magnitude and direction. This high degree of similarity suggests that it may be reasonable to directly interpret second- order associations found in the Fisher space.

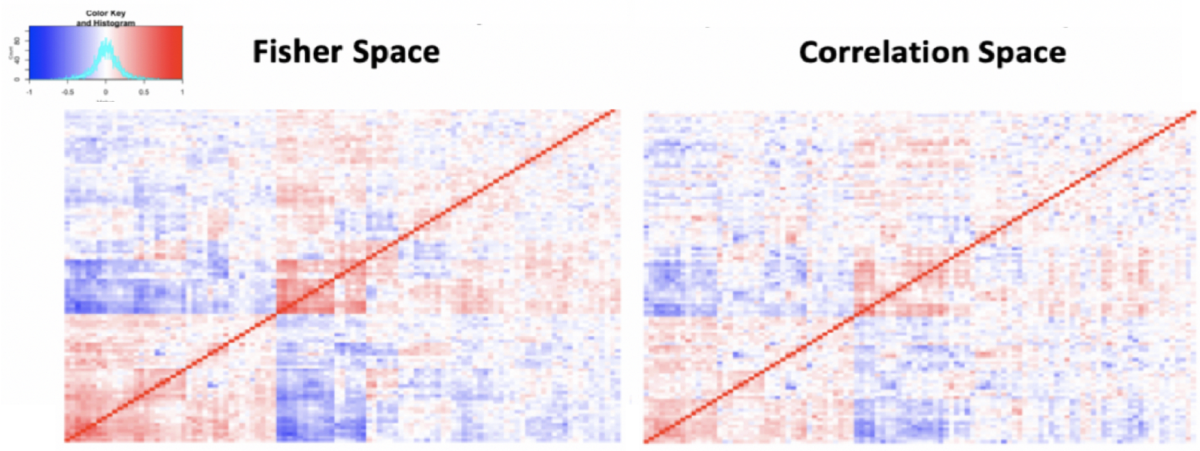


Figure 2: **Left Panel:** Heatmap of empirical second order correlation assesses inter-edge dependence in Fisher space. Rows and columns are clustered together based on similarity of correlation measure. **Right Panel:** Heatmap of empirical second order correlation assesses inter-edge dependence in the space of correlation networks. Row and column order are fixed to be the same as the heatmap assessing inter-edge dependence in Fisher space.

To supplement the results shown in section 4 of the main body of this manuscript, we now show stability selection results for every possible covariate-edge combination and every possible edge-edge second order association examined in our real data analysis. This information is visualized in the left panel of Figure 2 in the main body of this manuscript. Highlighted cells represent the edges flagged in the graph difference plots in Fisher space of Figure 3 in the main body of this manuscript. We note that edges displaying graph differences for a particular covariate all have higher stability selection thresholds than those edges found to have no graph differences for a particular covariate in Fisher space.

| Edge | ReadEng | PicVocab | Precuneus | Postcentral | Cuneus | Posteriorcingulate | Temporalpole |
|---|---|---|---|---|---|---|---|
| 1 | 0.005 | 0.005 | 0.115 | 0.005 | 0 | 0 | 0 |
| 2 | 0.02 | 0 | 0 | 0 | 0 | 0 | 0 |
| 3 | 0.005 | 0 | 0 | 0 | 0 | 0.025 | 0 |
| 4 | 0.005 | 0.005 | 0.125 | 0.26 | 0.07 | 0 | 0 |
| 5 | 0 | 0 | 0.13 | 0.185 | 0.02 | 0.01 | 0.005 |
| 6 | 0 | 0 | 0.005 | 0 | 0 | 0 | 0 |
| 7 | 0.005 | 0.01 | 0.015 | 0.01 | 0.005 | 0 | 0 |
| 8 | 0 | 0.005 | 0 | 0 | 0 | 0 | 0.01 |
| 9 | 0 | 0.045 | 0.26 | 0.065 | 0 | 0.02 | 0 |
| 10 | 0.02 | 0 | 0 | 0 | 0 | 0 | 0 |
| 11 | 0.025 | 0.04 | 0 | 0 | 0 | 0.005 | 0 |
| 12 | 0.14 | 0.06 | 0.005 | 0.035 | 0 | 0 | 0.005 |
| 13 | 0 | 0 | 0.025 | 0.005 | 0.305 | 0.01 | 0 |
| 14 | 0.005 | 0 | 0 | 0 | 0.01 | 0 | 0.015 |
| 15 | 0.05 | 0.15 | 0.16 | 0.44 | 0 | 0.02 | 0.01 |
| 16 | 0 | 0 | 0 | 0 | 0 | 0 | 0 |
| 17 | 0.005 | 0.19 | 0.02 | 0 | 0 | 0 | 0 |
| 18 | 0 | 0.01 | 0 | 0 | 0 | 0.005 | 0 |
| 19 | 0 | 0 | 0 | 0.005 | 0 | 0.005 | 0 |
| 20 | 0.5 | 0.27 | 0 | 0 | 0 | 0 | 0 |
| 21 | 0.105 | 0.415 | 0.105 | 0.055 | 0.075 | 0 | 0 |
| 22 | 0 | 0 | 0 | 0 | 0 | 0.095 | 0 |
| 23 | 0 | 0 | 0 | 0 | 0 | 0 | 0 |
| 24 | 0 | 0 | 0 | 0 | 0 | 0 | 0 |
| 25 | 0 | 0.01 | 0 | 0 | 0.03 | 0.045 | 0 |
| 26 | 0.02 | 0.015 | 0 | 0 | 0 | 0 | 0 |
| 27 | 0 | 0 | 0 | 0 | 0.005 | 0.02 | 0 |
| 28 | 0.015 | 0.005 | 0 | 0 | 0 | 0 | 0 |
| 29 | 0 | 0 | 0 | 0.005 | 0 | 0 | 0 |
| 30 | 0.005 | 0 | 0.05 | 0.895 | 0 | 0.04 | 0.005 |
| 31 | 0 | 0.01 | 0 | 0 | 0 | 0 | 0 |
| 32 | 0 | 0 | 0 | 0 | 0.005 | 0.005 | 0 |
| 33 | 0.05 | 0.115 | 0 | 0 | 0 | 0 | 0 |
| 34 | 0 | 0 | 0 | 0 | 0.105 | 0.01 | 0.005 |
| 35 | 0.055 | 0.095 | 0.01 | 0.12 | 0 | 0.015 | 0 |
| 36 | 0.025 | 0.01 | 0 | 0 | 0 | 0 | 0 |
| 37 | 0 | 0.005 | 0 | 0 | 0 | 0 | 0 |
| 38 | 0.005 | 0 | 0.005 | 0.025 | 0.01 | 0 | 0 |
| 39 | 0 | 0 | 0 | 0 | 0.025 | 0 | 0.005 |
| 40 | 0 | 0.01 | 0 | 0 | 0 | 0 | 0 |
| 41 | 0.005 | 0.005 | 0 | 0 | 0 | 0 | 0 |
| 42 | 0.39 | 0.275 | 0 | 0 | 0 | 0 | 0 |
| 43 | 0 | 0.02 | 0 | 0.035 | 0 | 0 | 0 |
| 44 | 0 | 0.005 | 0 | 0 | 0.005 | 0 | 0 |
| 45 | 0 | 0 | 0.825 | 0.015 | 0.015 | 0.015 | 0.005 |
| 46 | 0 | 0 | 0.01 | 0 | 0 | 0 | 0 |
| 47 | 0 | 0 | 0 | 0 | 0 | 0 | 0.005 |
| 48 | 0 | 0 | 0.57 | 0.105 | 0.005 | 0.11 | 0 |
| 49 | 0 | 0 | 0.15 | 0.195 | 0 | 0 | 0 |
| 50 | 0 | 0 | 0 | 0 | 0.015 | 0 | 0 |
| 51 | 0.005 | 0.01 | 0 | 0 | 0 | 0 | 0 |
| 52 | 0.005 | 0.005 | 0 | 0 | 0 | 0.005 | 0 |
| 53 | 0 | 0 | 0.005 | 0 | 0 | 0.015 | 0 |
| 54 | 0.005 | 0 | 0 | 0 | 0.01 | 0 | 0 |
| 55 | 0.005 | 0 | 0 | 0 | 0 | 0 | 0 |
| 56 | 0 | 0 | 0 | 0 | 0 | 0 | 0 |
| 57 | 0 | 0 | 0.01 | 0.13 | 0 | 0.095 | 0 |
| 58 | 0.025 | 0.005 | 0 | 0 | 0 | 0 | 0 |
| 59 | 0 | 0 | 0 | 0 | 0 | 0 | 0 |
| 60 | 0 | 0 | 0.72 | 0.125 | 0 | 0.01 | 0 |
| 61 | 0.005 | 0.14 | 0.01 | 0 | 0 | 0.005 | 0 |

| Edge | ReadEng | PicVocab | Precuneus | Postcentral | Cuneus | Posteriorcingulate | Temporalpole |
|---|---|---|---|---|---|---|---|
| 62 | 0 | 0 | 0 | 0 | 0 | 0 | 0 |
| 63 | 0 | 0 | 0 | 0 | 0.065 | 0 | 0.015 |
| 64 | 0 | 0.005 | 0.005 | 0.01 | 0 | 0 | 0 |
| 65 | 0.2 | 0.005 | 0 | 0 | 0 | 0 | 0 |
| 66 | 0 | 0 | 0 | 0 | 0 | 0 | 0 |
| 67 | 0 | 0 | 0.165 | 0 | 0 | 0 | 0 |
| 68 | 0 | 0 | 0 | 0 | 0 | 0 | 0 |
| 69 | 0 | 0 | 0 | 0 | 0 | 0 | 0 |
| 70 | 0 | 0 | 0 | 0 | 0 | 0 | 0 |
| 71 | 0 | 0 | 0 | 0 | 0 | 0 | 0 |
| 72 | 0 | 0.005 | 0.29 | 0 | 0 | 0 | 0 |
| 73 | 0.005 | 0.005 | 0.075 | 0.115 | 0 | 0 | 0.015 |
| 74 | 0 | 0 | 0.005 | 0 | 0.005 | 0 | 0.015 |
| 75 | 0 | 0 | 0.06 | 0.07 | 0 | 0 | 0 |
| 76 | 0.015 | 0.005 | 0 | 0 | 0 | 0 | 0.005 |
| 77 | 0 | 0.08 | 0.01 | 0 | 0 | 0 | 0 |
| 78 | 0 | 0 | 0.09 | 0.005 | 0.02 | 0.02 | 0.005 |
| 79 | 0 | 0 | 0 | 0 | 0 | 0 | 0 |
| 80 | 0 | 0.015 | 0.01 | 0 | 0 | 0 | 0 |
| 81 | 0 | 0 | 0 | 0 | 0 | 0 | 0 |
| 82 | 0.025 | 0 | 0.035 | 0.26 | 0.005 | 0.01 | 0.01 |
| 83 | 0.005 | 0 | 0.075 | 0 | 0.01 | 0.04 | 0 |
| 84 | 0 | 0 | 0 | 0.025 | 0.005 | 0.03 | 0 |
| 85 | 0.205 | 0.03 | 0 | 0 | 0 | 0 | 0 |
| 86 | 0.005 | 0.005 | 0.005 | 0 | 0 | 0 | 0 |
| 87 | 0 | 0 | 0.005 | 0.005 | 0 | 0 | 0.04 |
| 88 | 0.025 | 0.185 | 0 | 0 | 0 | 0 | 0 |
| 89 | 0 | 0 | 0 | 0 | 0.065 | 0 | 0.005 |
| 90 | 0.045 | 0.015 | 0.025 | 0.02 | 0.005 | 0 | 0 |
| 91 | 0.005 | 0.005 | 0.02 | 0 | 0 | 0.005 | 0 |
| 92 | 0 | 0 | 0.06 | 0.17 | 0.02 | 0 | 0.195 |
| 93 | 0.005 | 0.05 | 0.635 | 0.245 | 0 | 0.01 | 0 |
| 94 | 0.04 | 0 | 0.005 | 0 | 0 | 0 | 0 |
| 95 | 0 | 0 | 0 | 0 | 0 | 0 | 0 |
| 96 | 0.005 | 0.175 | 0 | 0.005 | 0 | 0 | 0 |
| 97 | 0 | 0 | 0.07 | 0.06 | 0 | 0 | 0.01 |
| 98 | 0 | 0 | 0 | 0 | 0 | 0 | 0 |
| 99 | 0 | 0.005 | 0 | 0.005 | 0 | 0 | 0.005 |
| 100 | 0 | 0 | 0 | 0 | 0.03 | 0 | 0 |
| 101 | 0.005 | 0.02 | 0 | 0 | 0 | 0 | 0.005 |
| 102 | 0 | 0.005 | 0 | 0.31 | 0.135 | 0 | 0.03 |
| 103 | 0 | 0 | 0 | 0 | 0 | 0 | 0.005 |
| 104 | 0 | 0 | 0.125 | 0 | 0 | 0 | 0.005 |
| 105 | 0 | 0 | 0 | 0 | 0 | 0 | 0 |

Table 6: Stability selection results for all potential covariate-edge associations over 200 bootstrap samples. Highlighted cells represent those displayed in graph difference plots in Figure 4

We now supplement Figure 4 with stability selection results for all potential second order edge-edge associations. We demonstrate, for each edge-edge association, nodes involved in each second-order pair, stability selection proportions over 200 bootstrap samples, and average partial correlation over 200 bootstrap samples. Highlighted rows represent those shown in Figure 4 and demonstrate the abundance of second-order associations in our dataset.

| Edge 1 | Edge 2 | Stability Selection | $\omega$ |
|---|---|---|---|
| (13, 8) | (13,10) | 1 | 0.863 |
| (8,1) | (11,8) | 1 | -0.740 |
| (8,3) | (13,3) | 1 | 0.716 |
| (8,1) | (13,1) | 1 | 0.657 |
| (9,1) | (13,9) | 1 | -0.634 |
| (9,1) | (10,9) | 1 | -0.591 |
| (12,8) | (13,8) | 1 | -0.578 |
| (5,3) | (9,3) | 1 | -0.533 |
| (6,3) | (15,3) | 1 | -0.517 |
| (8,3) | (10,3) | 1 | 0.517 |
| (12,8) | (13,12) | 1 | 0.509 |
| (8,1) | (8,3) | 1 | -0.495 |
| (8,1) | (10,1) | 1 | 0.495 |
| (9,1) | (11,1) | 1 | -0.487 |
| (11,8) | (13,11) | 1 | 0.478 |
| (8,4) | (13,4) | 1 | 0.467 |
| (11,1) | (11,8) | 1 | 0.465 |
| (11,1) | (13,11) | 1 | -0.449 |
| (13,1) | (13,11) | 1 | -0.449 |
| (13,6) | (15,13) | 1 | -0.436 |
| (11,1) | (12,1) | 0.995 | 0.433 |
| (12,3) | (13,3) | 1 | 0.433 |
| (11,2) | (11,6) | 1 | 0.431 |
| (12,6) | (15,12) | 1 | -0.407 |
| (13,11) | (15,13) | 1 | 0.406 |
| (10,8) | (13,8) | 1 | -0.406 |
| (6,1) | (15,1) | 1 | -0.391 |
| (3,1) | (4,3) | 1 | -0.388 |
| (5,4) | (9,4) | 1 | -0.388 |
| (11,1) | (11,6) | 1 | 0.387 |
| (8,1) | (9,8) | 1 | -0.378 |
| (12,1) | (12,8) | 1 | 0.375 |
| (7,6) | (13,7) | 1 | 0.366 |
| (9,8) | (11,8) | 1 | -0.363 |
| (11,3) | (15,3) | 1 | 0.358 |
| (4,1) | (8,4) | 1 | 0.357 |
| (9,4) | (13,4) | 0.99 | 0.352 |
| (8,3) | (12,8) | 0.995 | -0.349 |
| (14,8) | (14,13) | 1 | 0.349 |
| (8,1) | (10,8) | 0.965 | 0.346 |
| (3,1) | (8,3) | 0.995 | 0.344 |
| (13,1) | (13,8) | 0.935 | 0.343 |
| (13,1) | (13,6) | 1 | 0.343 |
| (11,9) | (13,4) | 1 | -0.343 |
| (5,4) | (9,1) | 0.99 | -0.338 |
| (12,11) | (15,12) | 1 | 0.335 |
| (11,6) | (15,11) | 1 | -0.334 |
| (4,2) | (6,4) | 1 | 0.334 |
| (8,4) | (9,8) | 0.995 | -0.333 |
| (13,4) | (13,8) | 0.94 | -0.333 |
| (8,5) | (10,8) | 1 | -0.332 |
| (9,3) | (11,3) | 1 | -0.33 |
| (13,9) | (13,10) | 1 | -0.327 |
| (12,11) | (13,12) | 1 | -0.322 |

| Edge 1 | Edge 2 | Stability Selection | $\omega$ |
|---|---|---|---|
| (2,1) | (6,1) | 1 | 0.320 |
| (5,4) | (15,4) | 1 | 0.317 |
| (9,8) | (13,9) | 0.99 | 0.311 |
| (3,2) | (6,3) | 1 | 0.310 |
| (13,3) | (13,12) | 0.995 | -0.307 |
| (10,9) | (14,10) | 1 | 0.307 |
| (8,3) | (14,3) | 1 | -0.307 |
| (15,8) | (15,13) | 1 | 0.305 |
| (9,1) | (9,4) | 0.985 | -0.305 |
| (9,1) | (10,5) | 0.995 | -0.304 |
| (12,11) | (15,11) | 1 | -0.303 |
| (7,5) | (7,6) | 1 | 0.302 |
| (6,4) | (15,4) | 1 | -0.302 |
| (11,5) | (15,11) | 1 | 0.301 |
| (4,1) | (9,4) | 0.995 | -0.301 |
| (12,10) | (15,12) | 0.995 | 0.3 |
| (9,5) | (11,9) | 1 | -0.299 |
| (11,8) | (12,1) | 0.82 | 0.299 |
| (8,1) | (8,4) | 0.955 | -0.294 |
| (11,1) | (13,8) | 0.98 | 0.293 |
| (8,6) | (15,8) | 1 | -0.291 |
| (11,10) | (13,9) | 0.995 | -0.291 |
| (13,2) | (13,9) | 0.985 | -0.29 |
| (8,1) | (12,3) | 0.985 | 0.286 |
| (12,1) | (12,6) | 0.99 | 0.285 |
| (9,1) | (14,10) | 0.995 | 0.284 |
| (7,6) | (14,6) | 1 | 0.282 |
| (9,2) | (12,6) | 1 | -0.278 |
| (10,8) | (11,9) | 0.985 | 0.278 |
| (12,1) | (13,4) | 1 | 0.278 |
| (13,4) | (14,10) | 1 | -0.277 |
| (11,8) | (13,8) | 0.975 | -0.276 |
| (13,5) | (13,10) | 1 | -0.275 |
| (12,1) | (15,8) | 0.955 | -0.275 |
| (14,1) | (14,4) | 1 | -0.274 |
| (11,6) | (12,8) | 0.99 | -0.273 |
| (6,5) | (13,5) | 1 | 0.273 |
| (12,8) | (15,8) | 0.88 | -0.272 |
| (11,9) | (15,11) | 1 | 0.272 |
| (10,5) | (15,11) | 0.995 | -0.272 |
| (13,11) | (13,12) | 0.995 | 0.272 |
| (3,1) | (13,3) | 0.99 | -0.27 |
| (10,8) | (11,10) | 0.97 | 0.268 |
| (9,1) | (12,6) | 0.945 | 0.265 |
| (8,3) | (9,8) | 0.98 | -0.265 |
| (10,4) | (11,10) | 1 | -0.264 |
| (4,1) | (11,4) | 1 | 0.262 |
| (9,6) | (11,2) | 1 | -0.262 |
| (4,2) | (9,2) | 1 | 0.260 |
| (8,3) | (15,3) | 1 | -0.257 |
| (6,1) | (15,6) | 0.995 | 0.256 |
| (5,1) | (13,5) | 1 | -0.256 |
| (12,8) | (13,3) | 0.685 | -0.255 |
| (13,8) | (13,9) | 0.965 | 0.252 |
| (12,9) | (12,10) | 0.99 | -0.252 |
| (10,3) | (13,3) | 0.995 | -0.251 |
| (12,1) | (13,1) | 0.9 | 0.251 |

Table 7: Stability selection results for all potential second order associations over 200 bootstrap samples. Columns represent, from left to right, nodes involved in each edge, stability selection proportion over 200 bootstrap samples, and average partial correlation between edges over 200 bootstrap samples.

We next provide a more comprehensive biological interpretation of flagged edges for covariates found to be statistically significant. These flagged covariate-dependent edges were demonstrated, via estimable differences, in Figure 3 of the main body of this manuscript.

**ReadEng:** ReadEng is a language processing covariate that involves a test in which participants are scored as correct or incorrect based on how accurately they read and pronounce letters and words. Flagged edges include the association between node 5 (Cerebellum and Amygdala) and node 7 (Cerebellum and Brain Stem) as well as the association between node 6 (Thalamus and Putamen) and node 10 (Cerebellum and Amygdala). The cerebellum is best known for its role in motor learning, maintaining balance, and the coordination of voluntary movements; recent studies indicate that the cerebellum also has a role in cognitive functions such as language recognition (Koziol et al. 2014). The brain stem controls autonomous processes (such as breathing and heart beat) and serves as a point to relay nerve information (Basinger and Hogg 2019). The amygdala plays a key role in processing memory, emotional responses (e.g. fear and anxiety), and decision making while the thalamus controls consciousness, relays sensory and motor signals, and is associated with episodic memory formulation and emotional expression (AbuHasan et al. 2020, Torrico and Munakomi 2019). Finally, the putamen is involved in learning and motor control and has also been linked to the reward and addiction process (Ghandili and Munakomi 2020). Given recent research implicating the cerebellum's role in language recognition, graph differences between network edges seem to link language recognition with emotional response and link language recognition with episodic memory and emotional expression.

**Precuneus Area:** The precuneus is involved in a variety of cognitive processes including recollection and memory, integration of environmental information, mental imagery strategies, recall of episodic memory, and affective responses to pain (Tomasi and Volkow 2020, Zhang and Chiang-shan 2012, Vogt and Laureys 2005, Berkovich-Ohana et al. 2020, Tanaka and Kirino 2016, Fauchon et al. 2019). In addition, the precuneus is a "hub" (highly interconnected) node in the default mode network, a functional connectivity subnetwork most associated with resting state (Utevsky et al. 2014). Flagged edges including the association between node 2 (Cerebellum and Brain Stem) and node 15 (Brain Stem and Hippocampus), the association between node 3 (Brain Stem and Pallidum) and node 11 (Cerebellum and Thalamus), the association between node 5 (Cerebellum and Amygdala) and node 12 (Thalamus and Diencephalon), the association between node 6 (Thalamus and Putamen) and node 13 (Putamen and Cerebellum), and the association between node 9 (Hippocampus and Cerebellum) and node 10 (Cerebellum and Amygdala). The pallidum plays a critical role in processing and executing motivated behaviors; it has a noted role in the process of addiction (Root et al. 2015). The diencephalon relays sensory information and controls autonomous functions of peripheral nervous system while the hippocampus plays major roles in memory and learning (Suryadevara et al. 2018, Anand and Dhikav 2012). Given precuneus function, edges seem to link memory and motor movement, motor movement and episodic memory, emotion and episodic memory, episodic memory and addiction, and memory and emotion.

**Postcentral Gyrus Area:** The postcentral gyrus is located in the primary somatosensory cortex and is the main receptor area for sense of touch (DiGuiseppi and Tadi 2020, Pijnenburg et al. 2015). In addition, the postcentral gyrus has been identified as a hub for somatosensory functional connectivity subnetworks (Tomasi and Volkow 2011, Fu et al. 2019). Flagged edges include the association between node 2 (Cerebellum and Brain Stem) and node 9 (Hippocampus and Cerebellum), node 2 (Cerebellum and Brain Stem) and node 15 (Brain Stem and Hippocampus), node 5 (Cerebellum and Amygdala) and node 6 (Thalamus and Putamen), and node 11 (Cerebellum and Thalamus) and node 15 (Brain Stem and Hippocampus). Given the known function of the postcentral gyrus, the majority of edges seem to link memory with motor movement, with the association between node 5 and node 6 possibly linking emotional response with episodic memory.

Given flagged covariate-dependent edges for statistically significant covariates, we conclude this section with a more comprehensive biological interpretation of key second order edge-edge associations displayed in Figure 4.

**Node 1 ↔ Node 8 | Node 8 ↔ Node 11:** This second order pair links the association between node 1 (Amygdala and Hippocampus) and node 8 (Cerebellum and Brain Stem) with the association between node 8 (Cerebellum and Brain Stem) and node 11 (Cerebellum and Thalamus). The amygdala and hippocampus (both part of the limbic system) overlap in function with the thalamus. The overlapping functions of these three areas (i.e. roles in memory) may contribute to the correlation between how nodes 1, 11 interact with node 8.

**Node 3 ↔ Node 8 | Node 3 ↔ Node 13:** This pair links the association between node 3 (Brain Stem and Pallidum) and node 8 (Cerebellum an Brain Stem) with the association between node 3 (Brain Stem and Pallidum) and node 13 (Putamen and Cerebellum). Along with the heavy overlap between anatomical areas, both edges associate the regulation of motor movements with reward and addiction.

**Node 8 ↔ Node 13 | Node 10 ↔ Node 13:** This pair links the association between node 8 (Cerebellum and Brain Stem) and node 13 (Putamen and Cerebellum) with the association between node 10 (Cerebellum and Amygdala) and node 13 (Putamen and Cerebellum). Similarly to the previous second-order pair, the link between motor control, emotion, and motivated behaviors (i.e. addiction) links the two edges together.
